# HNF1A is a novel BRD4 target and critical for BET-inhibitor response in pancreatic ductal adenocarcinoma

**DOI:** 10.1101/2025.10.01.679805

**Authors:** Kennedy S. Humphrey, Katherine J. Crawford, Bharani Muppavarapu, Melanie M. Mayberry, William Morris, Evan Torres, Mark D. Long, Jianxin Wang, Erik S. Knudsen, Agnieszka K. Witkiewicz, Ethan V. Abel

## Abstract

Pancreatic ductal adenocarcinoma (PDAC) is an exceedingly lethal cancer that lacks actionable molecular drivers, limiting precision treatment options. We previously identified the transcription factor, HNF1A, as a novel driver of tumorigenesis and pancreatic cancer stem cell (PCSC) properties in PDAC; however, HNF1A-targeting modalities do not currently exist. Here we show that HNF1A is a direct target of the epigenetic reader protein, BRD4, and its expression is exquisitely sensitive to BET-inhibitors (BETi), which inhibit PDAC cell proliferation and block PCSC-properties in a panel of HNF1A-expressing cell lines and patient-derived xenograft cells. Remarkably, we report that the antineoplastic activity of BETi/BRD4 knockdown can be overcome by restoration of HNF1A expression, but not by re-expression of canonical BETi target MYC. RNA-sequencing analyses revealed that a subset of BETi-responsive transcripts is dependent on HNF1A expression, including receptor tyrosine kinases (RTKs), regulators and ligands. Consistent with these data, we found that HNF1A restoration rescued EGFR/ERBB3-signaling and the protective effects of HNF1A restoration could be overcome with EGFR-inhibitors. Furthermore, we found that expressions of HNF1A, BRD4, and ERBB3 were strongly correlated across PDAC patient samples using multispectral immunofluorescence, supporting a connection between these players in PDAC biology, and high expression of ERBB3 associated with better survival, supporting the clinical importance of this network in patient outcomes. These findings demonstrate that BETi can be used to ablate HNF1A expression and that the inhibition of HNF1A is critical for BETi activity, while supporting HNF1A as novel therapeutic target in PDAC.

**Significance:** This study demonstrates that the oncogenic transcription factor HNF1A is a direct target of BRD4, and that the ablation of HNF1A by BET-inhibitors is central to their antineoplastic activity in PDAC.

## Introduction

Pancreatic ductal adenocarcinoma (PDAC) has a dismal 5-year survival rate of 13.3% and is projected to be the 2^nd^ leading cause of cancer related deaths by 2030^1^. In contrast to many other cancers where known oncogenes have led to effective targeted therapies, PDAC suffers from a dearth of targeted therapies despite an increasing comprehensive understanding of the biology of the disease. Therapeutic agents targeting KRAS, which is mutated in >90% of PDAC cases, are still in their infancy, while other targeted therapies such as MEK-inhibitors have underperformed for unclear reasons. Erlotinib, an EGFR-inhibitor, remains one of the few targeted therapies approved for PDAC^2, 3^, but its benefits are often modest, and it remains unclear what factors dictate response in the clinic. As such, there is a need to identify additional actionable targets and driver programs to expand treatment options and potentially predict response rates in PDAC patients.

In recent years there has been an expansion in the appreciation of transcription factors, epigenetic modulators, and transcriptional programs in PDAC biology. At least two dominant transcriptional subtypes (herein referred to as classical and non-classical) of PDAC have been identified by multiple groups, each being able to stratify patient survival. These subtypes have also been able to stratify responses to targeted therapies, such as erlotinib^3, 4^ and dasatinib^5^, and conventional chemotherapeutics^4, 5^, although the underlying determinants of these differing responses remain unclear. MYC, a well understood oncogenic transcription factor, has been shown to play roles in non-classical subtype determination and response to KRAS-pathway inhibition in PDAC^6, 7^. Other transcription factors, such as GATA6^8^ and TP63^8, 9^, have been implicated as master regulators of the classical and non-classical subtype, respectively, but their roles in therapeutic response and whether they are “actionable” drivers remains to be determined. Nonetheless, efforts to target transcription factors directly, or indirectly through targeting upstream and downstream epigenetic modulators, are increasingly being explored in PDAC.

Bromodomain and extra-terminal domain (BET) proteins are a family of epigenetic reader proteins consisting of four members: BRD2, BRD3, BRD4, and testis-specific BRDT. Containing two non-redundant bromodomains (BD) each^10^, BET proteins bind to acetylated lysines on histones and transcription factors, allowing them to coordinate the recruitment of other transcriptional components such as the RNA polymerase II complex and control the transcription of a subset of genes^11^. BET proteins are widely studied in cancer biology for their abilities to control the transcription of oncogenes such as MYC^12^ and BCL2^13^. As such, multiple generations of BET-inhibitors (BETi), which mimic acetylated lysines and bind competitively to either (BD1- or BD2-selective) or both (pan-BETi) BD to block BET protein-dependent transcription^11^. BETi show antineoplastic activity in many cancer types including PDAC^14–16^, often attributed to the transcriptional inhibition of MYC^12, 17–19^. However, the efficacy of BETi in PDAC does not strictly correspond to inhibition of MYC^15^, suggesting that other critical targets of BETi exist in PDAC.

Hepatocyte nuclear factor 1-alpha (HNF1A) is a gastrointestinal lineage transcription factor that has recently gained attention for driving malignancy programs in multiple cancers, including colorectal^20^, gastric^21^, and prostate cancers^22^. We reported that HNF1A is elevated in and a critical driver of pancreatic cancer stem cell (PCSCs)^23^ and metastasis^24^, both widely-pursued but currently untargetable components of PDAC biology. We found that knockdown of HNF1A inhibited proliferation and promoted apoptosis of PDAC cells, disrupted PCSC-properties such as tumorsphere formation and expression of the surface markers EPCAM, CD44 and CD24, and blocked tumor growth^23^, making HNF1A a potentially desirable target for PDAC. However, like many transcription factors, there are currently no direct means of targeting HNF1A expression or activity.

In this study, we explored whether BETi could be used to target HNF1A in PDAC in a manner analogous to MYC. We report that HNF1A expression is potently inhibited by multiple generations of BETi and is a direct transcriptional target of BRD4. Surprisingly, re-expression of HNF1A can overcome the antiproliferative effects of BETi and BRD4 knockdown both *in vitro* and *in vivo* and restores the expression of a subset BETi-responsive genes including EGFR-ligands and ERBB3. Consistent with this, we found that EGFR and ERBB3 phosphorylation were suppressed by BETi and rescued by HNF1A restoration, with EGFR inhibitors capable of overcoming HNF1A’s protective effects. These findings support that HNF1A is novel and critical target of BETi in PDAC and may shed light on the mechanisms behind transcriptional programs in PDAC and response to targeted therapeutics like erlotinib.

## MATERIALS AND METHODS

### Cell lines, cell culture, and inhibitors

AsPC-1(RRID: CVCL_0152), HPAF-II (RRID: CVCL_0313), and BxPC-3 (RRID: CVCL_0186) cells were purchased from ATCC (Manassas, VA). UM5, UM15 and UM53 (formally NY5, NY15, NY53, respectively) were established as low-passage cells isolated from human patient-derived xenografts as previously described^23^. PDX-derived cells were passaged less than 10 times in culture for these studies. All cells were maintained in RPMI-1640 with GlutaMAX™-I supplemented with 10% FBS (Gibco), 1% antibiotic-antimycotic (Gibco), and 100µg/ml gentamicin (Gibco). Cells were routinely tested for mycoplasma contamination using the MycoScope PCR Detection kit (Gentlantis, San Diego, CA) and authenticated by STR profiling (Roswell Park Comprehensive Cancer Center Genomics Shared Resource). All inhibitors were purchased from SelleckChem (Houston, TX).

### Lentiviruses

pENTR/D-TOPO/LacZ and pENTR/D-TOPO/HNF1A were previously described^23^. Human MYC was amplified from AsPC-1 cDNA with primers 5’-AGGCTCCGCGGCCGCCCCCTTCACCATGCCCCTCAACGTTAGCTTCACC-3’ and 5’-TGGGTCGGCGCGCCCACCCTTTTACGCACAAGAGTTCCGTAGCTG-3’ and cloned into pENTR/D-TOPO using NEBuilder HiFi DNA Assembly master mix and protocol (New England Biolabs). All cDNAs were shuttled into pLenti6.3/UbC/V5-DEST^23^ using LR Clonase II enzyme mix and protocol. miR30-based BRD4 targeting shRNAs (5’-AACGGTACCAAACACAACTCAATAGTGAAGCCACAGATGTATTGAGTTGTGTTTGGTACCGTG-3’ and 5’-AGACGAGATTGAAATCGACTTATAGTGAAGCCACAGATGTATAAGTCGATTTCAATCTCGTCG-3’) were purchased from Transomic Technologies but subcloned into pENTR/D-TOPO along with non-targeting shRNA (5’-ATCTCGCTTGGGCGAGAGTAAGTAGTGAAGCCACAGATGTACTTACTCTCGCCCAAGCGAGAG-3’). shRNAs were shuttled into pLentipuro5/TRE3G/V5-DEST, an all-in-one doxycycline inducible lentiviral vector developed for this study, using LR Clonase II enzyme mix and protocol. For the HNF1A promoter reporter, the HNF1A promoter was amplified with using primers 5’-GATATCCTCGAGAAGTCAAGGCTGCAGTAAGCCATGATCG-3’ and 5’- GGTGGCGGATCCGGCTCGGCTGCCACAGGGCCACGCGGCC-3’, digested with with XhoI and BamHI (New England Biolabs), and ligated upstream of PatGFP^23^ in pENTR/D-TOPO and recombined into pLentipuro3/BLOCK-iT-DEST^25^. Viruses were packaged in 293FT cells as previously described^23^. BRD4 shRNAs were co-packaged to maximize knockdown efficiency. Cells were treated with lentivirus-containing conditioned media for 72-96 hours and then selected with either blasticidin (for cDNAs) or puromycin (for shRNAs and HNF1A promoter reporter) for up to two weeks before use in studies. Pooled populations of cells overexpressing LacZ or HNF1A were continuously regenerated throughout the study to prevent genetic drift.

### Knockdown experiments

Non-targeting control (Ctl) (Cat# D-001810-01-20), HNF1A (Cat#M-008215-01-0005), MYC (Cat#M-003282-07-0005), BRD2 (Cat#M-004935-02-0005), BRD3 (Cat#M-004936-01-0005), and BRD4 (Cat#M-004937-02-0005) siRNAs were purchased from Dharmacon (Lafayette, CO) and were transfected at 25 nM using Lipofectamine® RNAiMAX Reagent (Invitrogen) for the times indicated in each figure legend.

### Western blot analysis

All lysates were collected and boiled in 1x Laemmli sample buffer with β-mercaptoethanol for 5 minutes followed by electrophoresis on 4-20% Mini-PROTEAN TGX precast Tris-Glycine-SDS gels (Bio-Rad, Hercules, CA), transfer to low-fluorescent PVDF (Bio-Rad) and incubated overnight in primary antibodies (1 µg/ml) in 1x Animal-Free Blocking Solution (Cell Signaling Technology, Danvers, MA) plus 0.1% Tween-20. Blots were incubated in DyLight™ 700 or 800-conjugated secondary antibodies in 5% milk in TBS plus 0.1% Tween-20 and 0.005% SDS at room temperature for 1 hour and imaged/quantitated by an Odyssey® CLx imaging system (Li-Cor, Lincoln, NE). β-Actin (AC-15; RRID: AB_10950489) antibody was purchased from Thermo Fisher Scientific. HNF1A (D7Z2Q; RRID: AB_2728751), HNF4A (C11F12; AB_2295208), HNF4G (E4V2B), MYC (D3N8F; RRID: AB_2631168), BRD2 (D89B4; RRID: AB_10835146), BRD4 (E1Y1P), phospho-RB (S807/811) (D20B12; RRID: AB_111786578), phospho-Histone H3 S10 (D7N8E; RRID: AB_2799431), and GFP (D5.1; RRID: AB_1196615) were purchased from Cell Signaling. BRD3 (2088C3a; RRID: AB_868478) antibody was purchased from Abcam (Cambridge, MA).

### Quantitative reverse transcription-PCR (qRT-PCR)

Total RNA was extracted using the Direct-zol RNA Miniprep kit and protocol (Zymo Research, Tustin, CA) and reverse transcribed with High-Capacity RNA-to-cDNA Master Mix (Applied Biosystem). SYBR Green PCR was performed using cDNA and iTaq Universal SYBR Green Supermix (Bio-Rad) on a QuantStudio 7 Pro Real-Time PCR System with QuantStudio Design & Analysis Software 2.8.0 (ThermoFisher Scientific). Conditions used for qPCR were 95°C for 10 minutes, followed by 40 cycles of 95°C for 10 seconds, 60°C for 15 seconds, and 72°C for 20 seconds. Primers for HNF1A were 5’-CCCACCAAGCAGGTCTTCAC-3’ and 5’-TGAAGGTCTCGATGACGCTG-3’. All quantitations were normalized to an endogenous control *ACTB* (5’-TACCTCATGAAGATCCTCACC-3’ and 5’-TTTCGTGGATGCCACAGGAC-3’). The comparative Ct method (ΔCt) was used to quantitate fold changes in HNF1A mRNA.

### Cell viability assay

Cells/colonies were incubated in 1x alamarBlue™ HS Cell Viability Reagent diluted in warmed complete media. Cells/colonies were allowed to incubate until reagent was ∼75% converted in one or more groups. Media was collected and analyzed for gain in fluorescence using 540nM excitation/590nM emission wavelengths on a BioTek Synergy HTX multi-mode reader (RRID: SCR_019749) and Gen5 3.03 software (RRID_ SCR_017317). All signal intensities were normalized back to their respective control groups, set to 100% viability.

### Colony formation assay

200 cells were plated per well of a 12-well cell culture plate. Cells were allowed to attach overnight before treatment with inhibitors. Media and inhibitors were replaced every three days for two to three weeks, depending on cell line, to allow for colony formation. At endpoint, colony viability was measured by alamarBlue™ assay as above, followed by fixing of colonies with 5% formaldehyde and staining with crystal violet for representative images. For knockdown studies, cells were transfected with siRNA 72 hours prior to plating for colony formation.

### Flow cytometry

For analysis of PCSC markers, cell labeling and analysis was performed as previously described^23^. Briefly, cells were trypsinized, washed with HBSS/2% FBS twice, and resuspended in HBSS/2% FBS at a concentration of 1 million cells/100µL. Primary antibodies were diluted 1:40 in cell suspensions and incubated for 30 minutes on ice with vortexing every 5 minutes. Cells were washed twice with HBSS/2% FBS and resuspended in this solution. Mouse anti-human EPCAM (CD326) clone HEA-125 (RRID: AB_2751108) was purchased from Miltenyi Biotec (San Diego, CA). Mouse anti-human CD44 clone G44-26 and CD24 clone ML5 purchased from BD Biosciences (San Jose, CA). Side scatter and forward scatter profiles were used to eliminate cell doublets. For PI staining, cells were trypsinized, washed in PBS and fixed in ice-cold 70% ethanol by drop-wise addition to cell suspensions and fixed at 4°C for 30 mins. Cells were then washed twice with PBS following removal of ethanol and RNA digestion was performed using 200µg/mL RNAase for 30 mins at 37°C. Cells were then stained with 167µg/mL propidium iodide for 30 mins. All flow cytometry experiments were performed using LSRII B analyzer (BD Biosciences, San Jose, CA; RRID: SCR_002159). Analyses for marker expression and cell cycle were performed using FCS Express 7 flow cytometry software (DeNovo Software, Pasadena, CA; RRID: SCR_016431).

### Tumorsphere assay

Cells grown under normal 2D conditions were either treated with inhibitors or siRNA for the time points indicated in the legends. Cells were then trysinized, washed, and resuspended as single cells in tumorsphere culture media containing 1% N2 supplement, 2% B27 supplement, 1% antibiotic-antimycotic, 20 ng/mL epidermal growth factor (Gibco, Carlsbad, CA), 20 ng/mL human bFGF-2 (Invitrogen), 10 ng/mL BMP4 (Sigma-Aldrich, St. Louis, MO), 10 ng/mL LIF (Sigma-Aldrich). Cells were plated in 12-well Ultra-Low Attachment Plates (Corning, Corning, NY), 200-1000 cells per well, depending on the cell line. Media with inhibitors or plain media (for knockdowns) was supplemented every 3 days. Tumorspheres were allowed to form for 10-14 days before manually counting.

### Mice and xenograft studies

NOD/SCID/IL2γR^-/-^ (NSG) were bred and maintained at Roswell Park Comprehensive Cancer Center’s animal care facilities. Evenly mixed sex mice, 8-10 weeks old were subcutaneously implanted with 250,000 tumor cells in a volume of 50 µl (1:1 volume of cell suspension in growth media and Matrigel) in the left midflank regions of the mice. Once tumors were measurable, mice were put on doxycycline (2mg/ml)/5% sucrose water and doxycycline chow (VWR) to induce the expression of the non-targeting or BRD4-targeting shRNAs. Tumor growth was monitored weekly by digital caliper and tumor volumes calculated by the (length x width^2^)/2 method, and all mice were sacrificed once tumors in any group reached 20mm^3^ in volume.

### Immunofluorescence staining of tissues

Multispectral immunofluorescent (mIF) staining on formalin-fixed paraffin-embedded (FFPE) PDAC tissue microarray (TMA) was performed using the Opal 6-Plex Detection Kit (AKOYA Biosciences, Marlborough, MA). HNF1A (GT4110, 1:150, Opal 620; RRID: AB_2538735) from ThermoFisher, BRD4 (E2A7X, 1:1000, Opal 780; RRID: AB_2687578), ERBB3 (D22C5, 1:75, Opal 570; RRID: AB_2721919), and phospho-ERBB3 Y1289 (D1B5, 1:200, Opal 690; RRID: AB_11178795) from Cell Signaling, FGFR4 (sc-136988, 1:75, Opal 520; RRID: AB_2103663) from Santa Cruz Biotechnology, and Cytokeratin AE1/AE3 (M3515, 1:150, Opal 480; RRID: AB_2132885) from Agilent Dako were used for staining. Slides were imaged on the PhenoImager™ HT (AKOYA Biosciences). Further analysis of the slides was performed using inForm® Software v2.6.0 (AKOYA Biosciences). 220 PDAC patient cores^26^ with sufficient live cells (threshold > 100 cells per core) were stained and quantitated. Phenoptr software (RRID: SCR_026104) was used to report the percentage of cells that were positive for each protein stained. For correlation of staining intensities and survival data, average whole cell staining intensity was measured for each core. For the violin plot, the Pearson correlation coefficients were calculated for each core first. Then the R values from each core are used to make the violinplot. 115 patient cores with associated survival data were used for survival analyses. Likely dead cells were excluded from both percent positivity and staining intensity analyses. TMA cores were obtained from PDAC patients who had been treated at Thomas Jefferson University Hospitals between the years 2002 and 2010 Institutional Review Board approval.

### Chromatin immunoprecipitation

AsPC-1 cells were fixed with 1% formaldehyde for 10 minutes (one confluent 15cm plate per precipitation). Mnase-sheared chromatin was generated using the SimpleChIP Plus Enzymatic Chromatin IP kit and protocol (Cell Signaling). BRD4 (E1Y1P) or normal IgG (Cell Signaling) antibodies were used for precipitation of chromatin. Quantitative PCR was performed using immunoprecipitated DNA and 2% chromatin input DNA a previously described^23^. Percent Input for immunoprecipitated DNA was calculated using the formula 2% x 2^(Ct^ ^2%^ ^Input^ ^Sample-Ct^ ^IP^ ^Sample)^. Primers for the HNF1A promoter (5’-TTGGCTAGTGGGGTTTTGG-3’and 5’-GGCTCGGCTGCCACAGGG-3’) were used, as were primers for the HNF1A intron 2 (5’-TGAGGTCTTGCCATGTTGCCC-3’ and 5’-GTTTCTGTTATAGCAGACACAG-3’) as a negative cis-locus control. MYOD (primers 5’-AGACTGCCAGCACTTTGCTATC-3’ and 5’-ATAGAAGTCGTCCGTTGTGGC-3’) was used as a non-BRD4 target gene control.

### RNA sequencing and transcriptomic analysis

The sequencing libraries were prepared from 200ng total RNA purified using the TruSeq Stranded Total RNA kit (Illumina Inc). Following manufacturer’s instructions, the first step depleted rRNA from total RNA. After ribosomal depletion, the remaining RNA was purified, fragmented and primed for cDNA synthesis. Fragmented RNA was then reverse transcribed into first strand cDNA using random primers. The next step removed the RNA template and synthesized a replacement strand, incorporating dUTP in place of dTTP to generate ds cDNA. AMPure XP beads (Beckman Coulter) were used to separate the ds cDNA from the second strand reaction mix resulting in blunt-ended cDNA. A single ‘A’ nucleotide was then added to the 3’ ends of the blunt fragments. Multiple indexing adapters, containing a single ‘T’ nucleotide on the 3’ end of the adapter, were ligated to the ends of the ds cDNA, preparing them for hybridization onto a flow cell. Adapter-ligated libraries were amplified by PCR, purified using Ampure XP beads, and validated for appropriate size on a 4200 TapeStation D1000 Screentape (Agilent Technologies, Inc.). The DNA libraries were quantitated using KAPA Biosystems qPCR kit, and were pooled together in an equimolar fashion, following experimental design criteria.

Each pool was denatured and diluted to 16 pM for On-Board Cluster Generation and sequencing on a HiSeq2500 sequencer using the appropriate paired-end cluster kit and rapid mode SBS reagents following the manufacturer’s recommended protocol (Illumina Inc.). Raw feature counts were normalized and differential expression analysis using DESeq2. Differential expression rank order was utilized for subsequent Gene Set Enrichment Analysis (GSEA), performed using the fgsea (v1.8.0) package in R. Gene sets queried included those from the Hallmark Gene Sets available through the Molecular Signatures Database (MSigDB). Sample level gene set enrichment on normalized expression counts was performed via gene set variation analysis (GSVA)^27^, utilizing PDAC subtype gene sets^4, 28, 29^. Venn diagrams for overlapping gene sets were generated using DeepVenn (www.deepvenn.com).

### Statistical analysis

Data throughout are expressed as the mean ± SEM. Statistically significant differences between two groups were determined by the two-sided Student’s t-test for continuous data, while ANOVA was used for comparisons among multiple groups with Tukey’s or Dunnett’s post hoc tests where appropriate, with significance defined as p < 0.05. GraphPad Prism 10 was used for these analyses.

### Study approval

All animal protocols were approved by The Institute Animal Care and Use Committee (IACUC) at Roswell Park Comprehensive Cancer Center. The animal welfare assurance number for this study is A3143-01.

## RESULTS

### BET-inhibitors block HNF1A expression in PDAC cells

As BET-inhibitors (BETi) have been used extensively to block the expression of oncogenic transcription factors like MYC, we sought to determine whether this approach could be used to block HNF1A expression. Using conventional human PDAC cell lines AsPC-1 and HPAF-II, and human patient-derived xenograft (PDX) lines UM5 and UM15, all of which express HNF1A, we found that HNF1A expression was potently reduced by pan-BETi OTX-015, with partial reduction noted at 50nM and complete ablation at 500nM at 72 hours across all lines (**Fig. 1A**). Interestingly, MYC, which is most commonly associated with BETi, only showed partial reduction in expression by BETi in these cells, and only at higher concentrations of OTX-015. To better characterize the kinetics with which OTX-015 blocks HNF1A expression, we performed a time course study using both short (3 and 6 hours) and long (24, 48, and 72 hours) timepoints. HNF1A protein was mostly depleted by 24 hours in all cell lines, with decrease of the protein levels of HNF1A target genes HNF4A and HNF4G peaking between 48 and 72 hours (**Fig. 1B**).

**Figure 1.**
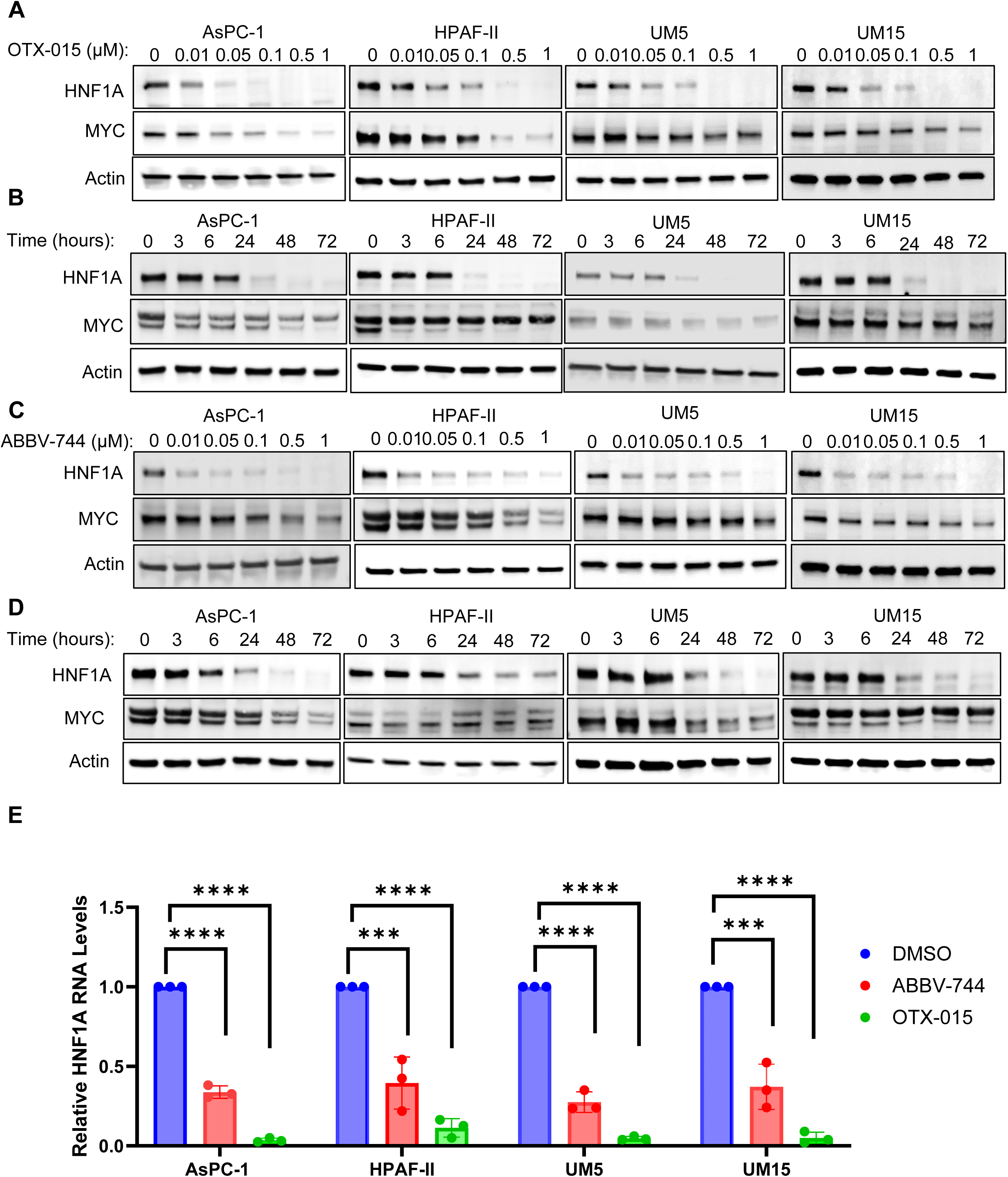
BETi block HNF1A expression in PDAC cells. **A,** Western blotting of HNF1A, MYC and Actin for AsPC-1, HPAF-II, UM5, and UM15 cells treated with the indicated doses of OTX-015 for 72 hours. **B,** Western blotting of HNF1A, MYC and Actin for AsPC-1, HPAF-II, UM5, and UM15 cells treated with 0.5 µM OTX-015 for the indicated time points. **C,** Western blotting of HNF1A, MYC and Actin for AsPC-1, HPAF-II, UM5, and UM15 cells treated with the indicated doses of ABBV-744 for 72 hours. **D,** Western blotting of HNF1A, MYC and Actin for AsPC-1, HPAF-II, UM5, and UM15 cells treated with 0.1 µM ABBV-744 for the indicated time points. **E,** quantitative RT-PCR analysis of HNF1A mRNA levels following 24 hours of treatment of AsPC-1, HPAF-II, UM5, and UM15 cells with either DMSO, 0.1 µM ABBV-744, or 0.5 µM OTX-015. Bar graphs represent the mean ± SEM, n=3. Statistical difference was determined by two-sided Student t-test with Welch’s correction; ns = non-significant, *p<0.05, **p<0.01, ***p<0.001, ****p<0.0001.

BET family proteins possess two bromodomains (BD1 and BD2) with overlapping as well as non-redundant activities. A number of bromodomain-selective BETi have been developed in recent years, with BD1-selective inhibitors showing similar activities to pan-BETi, while BD2-selective inhibitors affect the transcription of a subset of BET protein target genes, possibly resulting in a safer drug profile in patients. To determine whether BD-selective BETi have differential effects on the expression of HNF1A, we utilized the BD1-selective inhibitor GSK778 and BD2-selective inhibitors GSK046 and GSK620. While GSK778 most drastically reduced the expression of HNF1A in AsPC-1 cells, expression of HNF1A was also substantially diminished by GSK046 and GSK620 (Supp. Fig. 1A), indicating that the activity of both bromodomains of

BET proteins are required for HNF1A expression in PDAC cells. A third BD2-selective inhibitor, ABBV-744, also demonstrated the ability to reduce HNF1A expression at doses a low as 10nM in multiple PDAC lines (AsPC-1, HPAF-II, UM5 and UM15) (**Fig. 1C**) and as early as 24hrs (**Fig. 1D**), similar to OTX-015. Because of their high potencies and representation of distinct classes of BETi, we opted to use OTX-015 and ABBV-744 for the duration of this study. Consistent with transcriptional blocking of HNF1A expression, we found that both OTX-015 and ABBV-744 significantly reduced *HNF1A* mRNA levels within 24 hours of treatment (**Fig. 1E**).

### Inhibition of HNF1A expression is critical for BET-inhibitor activity

Stable transcriptional inhibition of oncogenic transcription factors and networks, such as MYC, have been shown to be critical for the responsiveness of cancer cells to BETi, and the restoration of these transcriptional programs can portend intrinsic or adaptive resistance to BETi^30^. Consistent with our previous studies^23, 24^, knockdown of HNF1A suppressed colony formation of PDAC cells (Supp. Fig. 1B), supporting its role as a critical oncogenic transcription factor. To determine whether restoration of HNF1A could similarly rescue the effects of BETi in PDAC, we overexpressed HNF1A from a BETi insensitive promoter (human UbC). Consistent with the HNF1A transcriptional network being disrupted by BETi, the expression of HNF1A target proteins HNF4A and HNF4G were reduced in response to BETi across multiple cell lines (**Fig. 2A**). Re-expression of HNF1A resulted in rescued expression of HNF4A and HNF4G in all cases (**Fig. 2A**). Additionally, we observed the rescued expression of MYC (**Fig. 2A**), and both the inhibition and rescue of HNF4A, HNF4G, and MYC occurred in dose-dependent manner mirroring inhibition/re-expression of HNF1A (**Fig. 2B**). The rescued expression of MYC in particular suggested that the antiproliferative effects of BETi may be dependent on HNF1A expression. In support of this, we observed that re-expression of HNF1A rescued cell viability across cell lines treated acutely (7 days) with either OTX-015 or ABBV-744 (**Fig. 2C**). To determine if rescue of MYC expression was responsible for the rescued viability observed with

**Figure 2.**
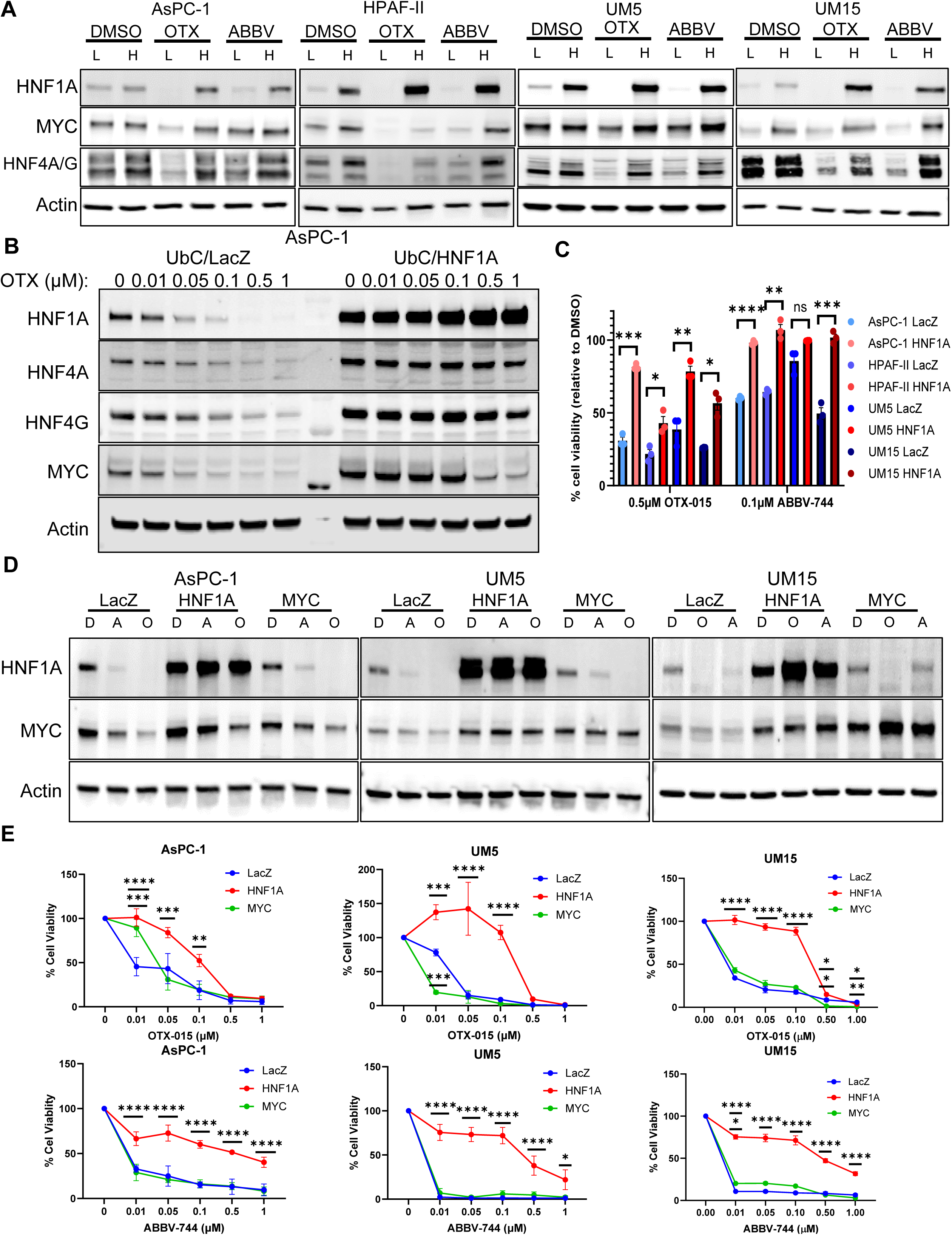
Effects of HNF1A and MYC re-expression on to BETi. **A,** Western blotting for HNF1A, HNF4A (top band), HNF4G (bottom band), MYC and Actin in LacZ or HNF1A overexpressing AsPC-1, HPAF-II, UM5, and UM15 cells treated with DMSO, 0.5 µM OTX-015, or 0.1 µM ABBV-744 for 72 hours to confirm overexpression and resistance to BETi. **B,** quantitation of LacZ or HNF1A overexpressing AsPC-1, HPAF-II, UM5, and UM15 cell viability (AlamarBlue) following 7 days of treatment with DMSO, 0.5 µM OTX-015, or 0.1 µM ABBV-744. Drugs were replenished every 3 days. Bar graphs represent the mean ± SEM, n=3. Statistical difference was determined by two-sided Student t-test with Welch’s correction; ns = non-significant, *p<0.05, **p<0.01, and ***p<0.001. **C,** Western blotting of HNF1A, MYC, HNF4A, HNF4G, and Actin in LacZ and HNF1A overexpressing AsPC-1 cells treated with the indicated doses of OTX-015. **D,** Western blotting for HNF1A, MYC and Actin in LacZ, HNF1A, and MYC overexpressing AsPC-1, UM5, and UM15 cells treated with DMSO, 0.5 µM OTX-015, or 0.1 µM ABBV-744 for 72 hours to confirm overexpression and resistance to BETi. **E,** dose curves colony formation experiments for LacZ, HNF1A, and MYC overexpressing AsPC-1, UM5, and UM15 cells. Cells were plated at 200 cells per well and treated with the indicated concentrations of OTX-015 or ABBV-744 for 2 weeks. Resultant colonies were quantitated by AlamarBlue viability assay. Data points represent the mean ± SEM, n=3. Statistical difference was determined by one-way ANOVA with Tukey’s multiple comparisons test; ns = non-significant, *p<0.05, **p<0.01, ***p<0.001, ****p<0.0001.

HNF1A re-expression, we overexpressed MYC using the human UbC promoter. Levels of re-expressed MYC were comparable to those observed with HNF1A re-expression in the presence of BETi (**Fig. 2D**). Interestingly, while HNF1A re-expression was able to rescue colony formation and viability across a range of BETi doses (**Fig. 2E, Supp.** Fig. 2A,B), re-expression of MYC was unable to rescue growth (**Fig. 2E**, Supp. Fig. 2B). Nonetheless, HNF1A overexpression could not rescue the anti-proliferative effects of direct MYC knockdown (Supp. Fig. 2C,D), indicating that while MYC expression is required for PDAC cell proliferation even with HNF1A present, its re-expression is not sufficient to rescue PDAC cell growth. Collectively, these data support that HNF1A re-expression drives resistance to BETi through processes beyond control of MYC expression.

We have previously shown that HNF1A contributes to both proliferation and stemness in PDAC^23, 24^. As such, we next sought to determine whether re-expression of HNF1A rescues cell cycle progression. At the protein level, we found that markers of cell cycle progression, including phosphorylated RB and Histone H3, were reduced by BETi treatment but rescued by HNF1A re-expression (**Fig. 3A**). Using propidium iodide (PI) staining, we found that BETi induced a G0/G1 arrest in AsPC-1, HPAF-II, and UM5 cells that could be rescued by HNF1A re-expression (**Fig 3B**, Supp. Fig 2E). Corresponding alterations in S phase and G2/M were partially or fully rescued by HNF1A re-expression in a cell-specific manner (**Fig 3B**, Supp. Fig 2E). We previously demonstrated that co-expression of the surface markers EPCAM (ESA) and CD44 identifies pancreatic cancer stem cells (PCSCs) and is responsive to HNF1A manipulation^23^. In line with this, we found that BETi reduced the co-expression of EPCAM and CD44, primarily through loss of EPCAM (an HNF1A target gene) expression (**Fig. 3C,D**), and the depletion of EPCAM+/CD44+ cells was reversed by re-expression of HNF1A. To determine if PCSC function is also disrupted in an HNF1A-dependent manner, we performed tumorsphere assays with AsPC-1 and HPAF-II cells treated for 7 days prior to replating in tumorsphere conditions and then either left untreated (pretreated) or maintained in continuous treatment of BETi. In both pretreated and continuously treated conditions, BETi significantly reduced tumorsphere formation, which in both cases were reversed by HNF1A re-expression (**Fig. 3F, G**). These findings suggest that PDAC stemness can be disrupted by either transient or continuous treatment with BETi, and that this stemness is dependent on HNF1A expression. Collectively, these data demonstrate that BETi prevent cell cycle progression and disrupt stemness in an HNF1A-dependent manner.

**Figure 3.**
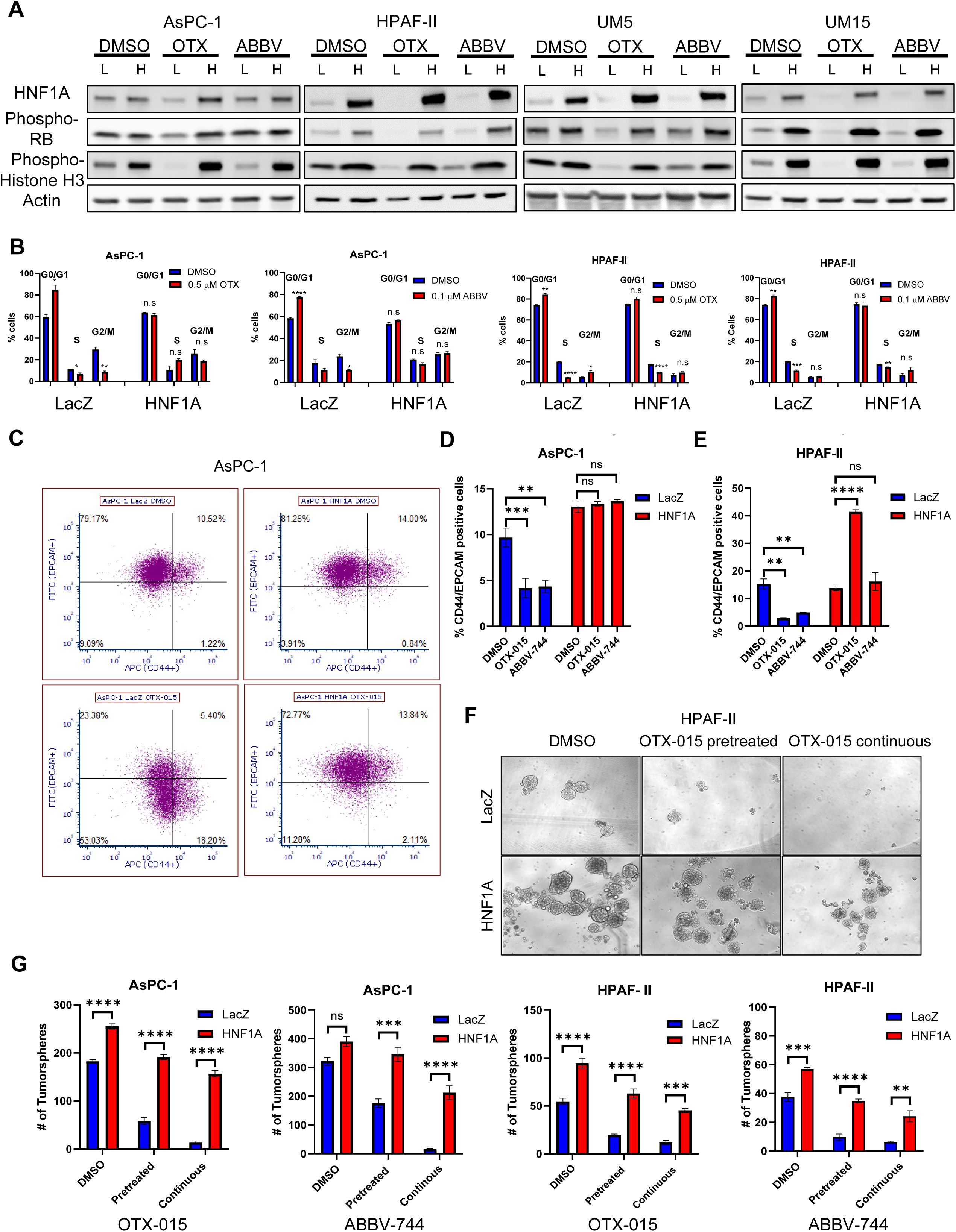
Re-expression of HNF1A restores cell cycle progression and stemness. **A,** Western blotting for HNF1A, phospho-RB S807/S811, phospho-Histone H3 S10, and Actin for LacZ and HNF1A overexpressing cells treated with 0.5 µM OTX-015 or 0.1 µM ABBV-744 for 72 hours. **B,** quantitation of propidium iodide staining of LacZ and HNF1A overexpressing AsPC-1 and HPAF-II cells treated with 0.5 µM OTX-015 or 0.1 µM ABBV-744 for 72 hours. Relative percentages of G0/G1, S, and G2/M phase cells are shown. **C,** flow cytometry analysis of CD44 and EPCAM surface expression on LacZ and HNF1A overexpressing AsPC-1 cells treated with 0.5 µM OTX-015 for 72 hours. **D,** quantitation of EPCAM+/CD44+ LacZ and HNF1A overexpressing AsPC-1 and **E,** HPAF-II cells treated with 0.5 µM OTX-015 or 0.1 µM ABBV-744 for 72 hours. **F,** representative images of tumorspheres formed from LacZ and HNF1A overexpressing HPAF-II cells. Cells were treated for one week in 2D cell culture with either DMSO or 0.5 µM OTX-015 before being replated in non-adherent tumorsphere culture. For OTX-015 treated cells, replated cells were either maintained in DMSO (pretreated) or additional 0.5 µM OTX-015 (continuous). Media and inhibitor were replenished every 3 days for 14 days before imagining and quantitation of spheres. **G,** LacZ and HNF1A overexpressing AsPC-1 and HPAF-II were treated with DMSO, 0.5 µM OTX-015 or 0.1 µM ABBV-744 as described in **F**. quantitation of resultant tumorspheres is shown. All bar graphs in this figure represent the mean ± SEM, n=3. Statistical difference was determined by one-way ANOVA with Tukey’s multiple comparisons test; ns = non-significant, *p<0.05, **p<0.01, ***p<0.001, ****p<0.0001.

### BRD4 directly regulates HNF1A expression in PDAC cells

The BET-family consists of four family members – the ubiquitously expressed BRD2, BRD3, and BRD4, and testes-specific BRDT – all of which are equivalently targeted by BETi. To determine which BET family members regulate HNF1A expression, we knocked down BRD2, BRD3, BRD4, or all three expressed proteins (BRDT was not expressed in any cells examined, data not shown). Knockdown of BRD4 reduced HNF1A protein and mRNA levels in cells tested, while knockdown of BRD2 also reduced HNF1A expression in AsPC-1 and UM15 cells (**Fig. 4A, B**).

**Figure 4.**
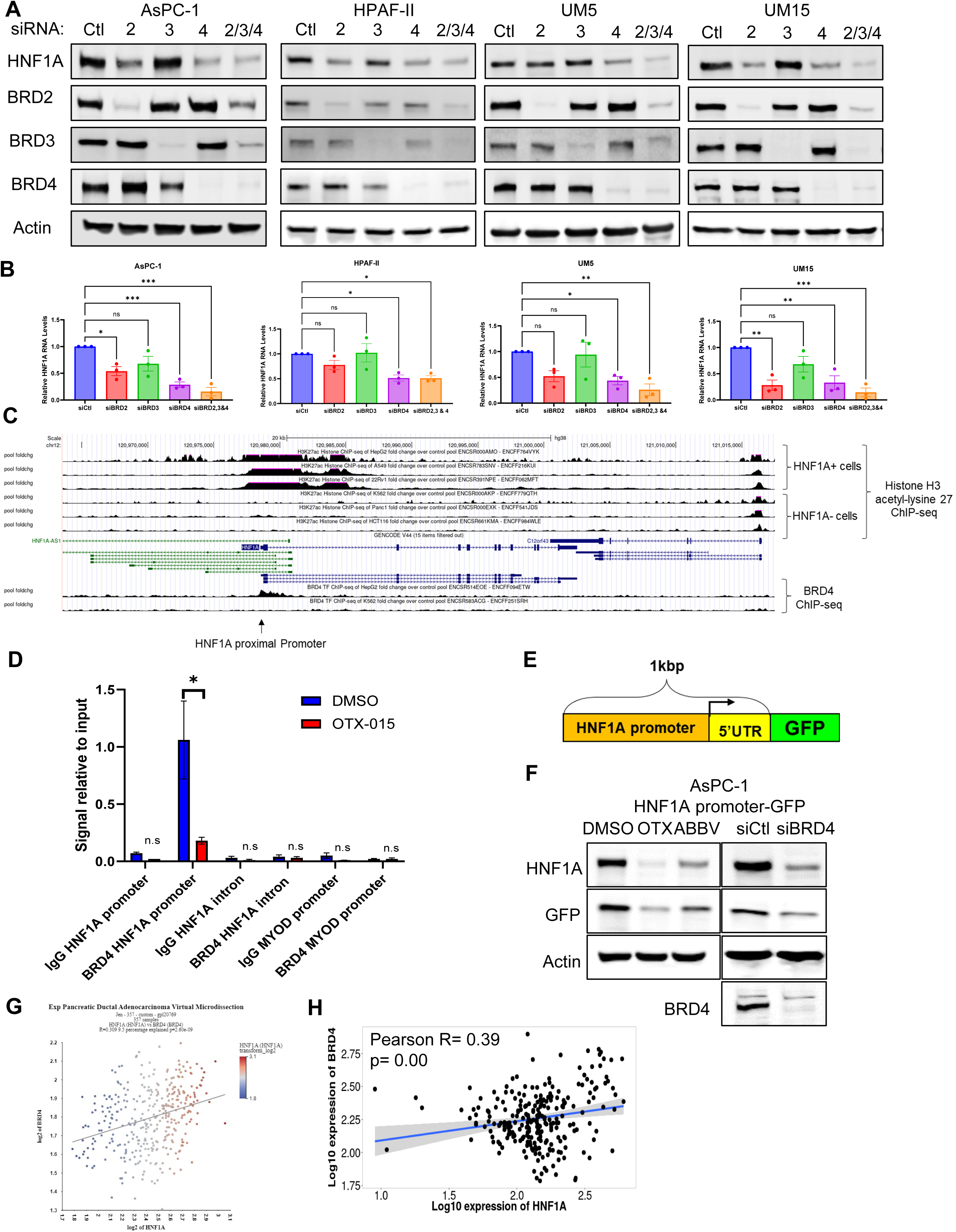
BRD4 directly regulates HNF1A expression. **A,** Western blotting for HNF1A, BRD2, BRD3, BRD4, and Actin for AsPC-1, HPAF-II, UM5, and UM15 cells transfected with control (Ctl), BRD2, BRD3, BRD4, or combined BET protein siRNAs for 96 hours. **B,** quantitative RT-PCR analysis of HNF1A mRNA levels following 96 hours of knockdown of AsPC-1, HPAF-II, UM5, and UM15 cells with Ctl or BRD4 siRNA. Bar graphs represent the mean ± SEM, n=3. Statistical difference was determined by two-sided Student t-test with Welch’s correction; ns = non-significant, *p<0.05, **p<0.01, ***p<0.001, ****p<0.0001. **C,** UCSC Genome Browser ChIP-seq tracks for the HNF1A locus. ChIP-seq experiments for Histone H3 acetyl-lysine 27 and BRD4 are shown for HNF1A-positive (HepG2, A594, 22Rv1) and HNF1A-negative (K562, Panc1, HCT116) cells. The HNF1A proximal promoter region is indicated. **D,** ChIP-PCR was performed on AsPC-1 cells treated with DMSO or 0.5 µM OTX-015 for 24 hours using normal IgG or BRD4 antibody. Enrichment of the HNF1A promoter was compared to the second intron of HNF1A as an intragenic control and the MYOD promoter as an inactive promoter control. Bar graphs represent the mean ± SEM, n=3. Statistical difference was determined by two-sided Student t-test with Welch’s correction; ns = non-significant, *p<0.05. **E,** schematic representation of a HNF1A reporter construct. A 1kbp region of the proximal promoter region, including the 5’ UTR was cloned upstream of a GFP reporter. **F,** Western blotting of HNF1A, GFP, BRD4, and Actin for AsPC-1 reporter cells treated with DMSO, 0.5 µM OTX-015 or 0.1 µM ABBV-744 for 24 hours or transfected with Ctl or BRD4 siRNA for 96 hours. **G,** correlation of HNF1A and BRD4 mRNA from Moffitt *et al*., 2015^29^. **H,** correlation of HNF1A and BRD4 mRNA from PDAC tumors from TNMplot.com^33^.

Additionally, treatment of AsPC-1 cells with the BRD4-selective degrader ZXH-3-26^31^ markedly reduced HNF1A expression without affecting BRD2 or BRD3 levels (Supp. Fig. 3A). The effects of combined knockdown of BET proteins on HNF1A expression were not significantly different from BRD4 knockdown alone, suggesting that BRD4 is the BET protein most responsible for regulation of HNF1A (**Fig. 4A, B**). To determine whether the regulation of HNF1A by BRD4 is direct, we mined publicly available BRD4 and Histone H3 acetyl-lysine 27 (a BRD4 binding moiety^10^) ChIP-sequencing (ChIP-seq) data from the ENCODE Project^32^. We found that Histone H3 acetyl-lysine 27 was enriched at the proximal promoter and first intron of HNF1A in HNF1A+ cells (HepG2, A549, and 22Rv1) but absent in HNF1A-cells (K562, Panc1, and HCT116) (**Fig. 4C**). Similarly, a broad BRD4 peak was present in HepG2 cells at the proximal promoter and the 5’ end of the first intron of HNF1A (**Fig. 4C**), supporting direct regulation of HNF1A by BRD4. To validate these findings, we performed ChIP-PCR in AsPC-1 cells treated with OTX-015 for 24 hours and found that BRD4 was both present at the HNF1A promoter and dissociated by treatment with BETi (**Fig. 4D**). We further validated the direct regulation of HNF1A by BRD4 by performing reporter assays using an HNF1A promoter-driven GFP lentiviral construct (**Fig. 4E**). We found that GFP expression was decreased in response to both BETi treatment and knockdown of BRD4 (**Fig. 4F**), indicating that BRD4 directly regulates transcription of HNF1A from its proximal promoter. Lastly, we found that mRNA levels of *HNF1A* and *BRD4* were significantly correlated in RNA-sequencing (RNA-seq) datasets from Moffitt *et al*., 2015^29^ and TNMplot^33^ (**Fig. 4G, H**), while correlation of HNF1A with BRD2 or BRD3 was either weaker or non-existent (Supp. Fig. 3B, C). Collectively, these data support that BRD4 is a direct regulator of HNF1A expression, and BETi directly block HNF1A transcription by dissociating BRD4 from its promoter.

To determine if genetic ablation of BRD4 phenocopies the biological effects of BETi in regard to HNF1A, we performed BRD4 knockdowns in our LacZ and HNF1A overexpressing PDAC lines. Similar to our earlier findings, the knockdown of BRD4 reduced the expression of HNF4A and HNF4G, which could be partially rescued by HNF1A re-expression (**Fig. 5A**). Re-expression of HNF1A was also able to rescue cell viability with short-term (96 hours) BRD4 knockdown (**Fig. 5B**) and in colony formation assays (**Fig. 5C, D**). As with BETi treatment, BRD4 knockdown depleted EPCAM+/CD44+ cells and tumorsphere formation, which were both rescued by HNF1A re-expression (**Fig. 5A, F**). Finally, to test the effects of BRD4 knockdown and HNF1A re-expression on tumor growth, we generated AsPC-1 LacZ and HNF1A cells with inducible BRD4 shRNA and implanted them subcutaneously into NSG mice. Once tumors were established, mice were administered doxycycline to induce the expression of the BRD4 shRNA. As expected, the knockdown of BRD4 greatly delayed tumor growth, while HNF1A re-expressing tumors grew at rates comparable to their control shRNA cohorts (**Fig. 5G**). Analyses of the resultant tumors showed that BRD4 knockdown was maintained throughout the duration of the experiment (**Fig. 5H**), while staining of tumors for Ki-67 revealed a reduction in proliferating cells in BRD4 knockdown tumors that were rescued by HNF1A re-expression (**Fig. 5I**).

**Figure 5.**
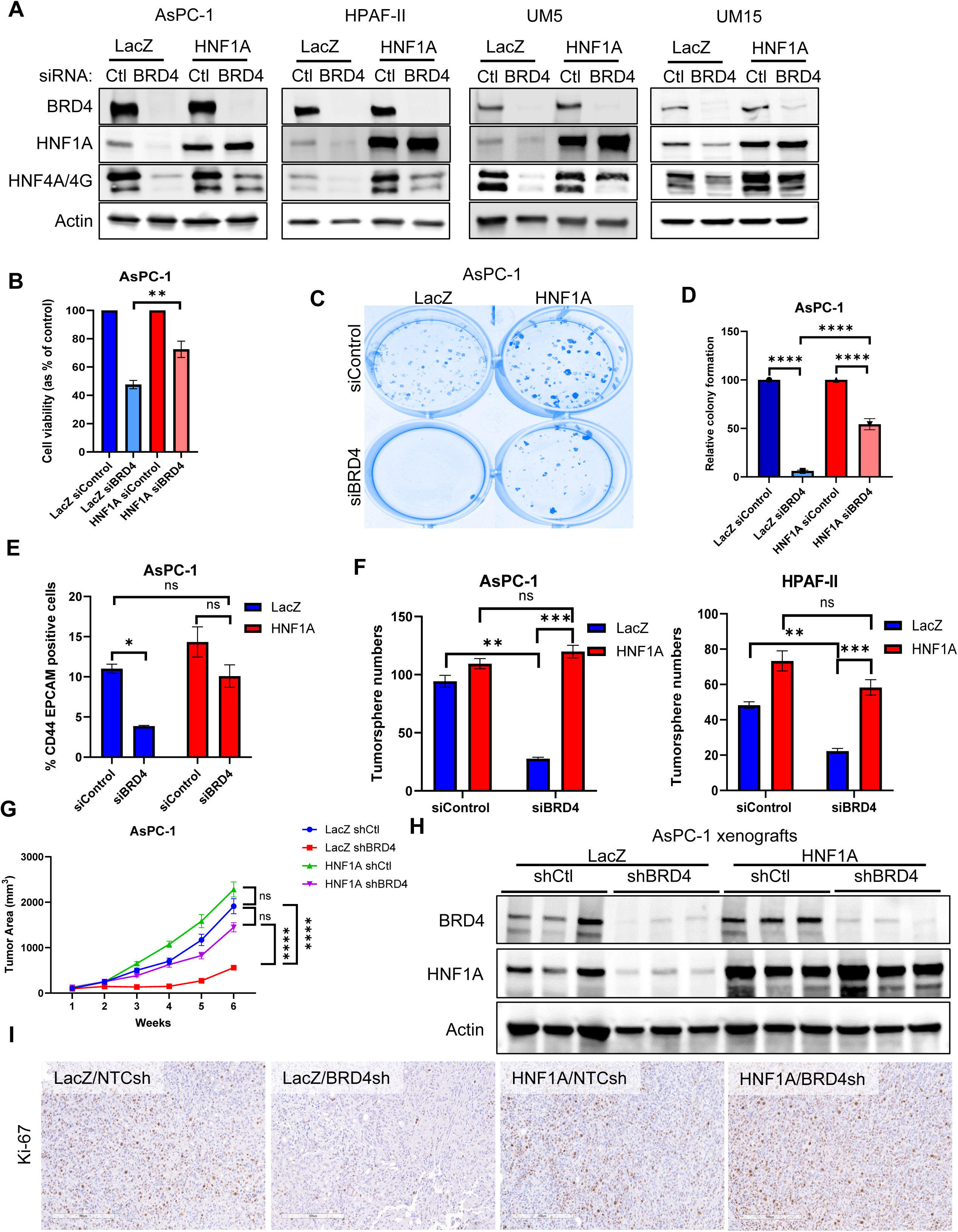
Re-expression of HNF1A overcomes effects of BRD4 knockdown. **A,** Western blotting of LacZ and HNF1A overexpressing cells transfected with Control (Ctl) or BRD4 siRNA for 96 hours for BRD4, HNF4A and HNF4G, HNF1A, and Actin. **B,** quantitation of cell viability of LacZ and HNF1A overexpressing AsPC-1 cells following 96 hours of BRD4 knockdown. Bar graphs represent the mean ± SEM, n=3. Statistical difference was determined by one-way ANOVA with Tukey’s multiple comparisons test; **p<0.01. **C,** representative AsPC-1 colonies after 2 weeks of BRD4 knockdown. siRNA was re-transfected after the first week. **D,** quantitation of colony viability performed in **C**. Bar graphs represent the mean ± SEM, n=3. Statistical difference was determined by one-way ANOVA with Tukey’s multiple comparisons test; ****p<0.0001. **E,** flow cytometry analysis of CD44 and EPCAM surface expression on LacZ and HNF1A overexpressing AsPC-1 cells following 96 hours of BRD4 knockdown. Bar graphs represent the mean ± SEM, n=3. Statistical difference was determined by one-way ANOVA with Tukey’s multiple comparisons test; ns = non-significant, *p<0.05. **F,** tumorsphere formation of LacZ and HNF1A overexpressing AsPC-1 and HPAF-II cells. Cells were transfected 72 hours prior to replating and grown under tumorsphere conditions for 2 weeks. Bar graphs represent the mean ± SEM, n=3. Statistical difference was determined by one-way ANOVA with Tukey’s multiple comparisons test; ns = non-significant, **p<0.01, ***p<0.001. **G,** subcutaneous tumor growth of LacZ and HNF1A overexpressing AsPC-1 cells. Once tumors reached a measurable size, mice were administered doxycycline chow and water to induce control or BRD4 shRNA expression. Tumors were monitored until humane endpoint was reached in any group. Data points represent the mean tumor volumes ± SEM. n=11 mice per group. Statistical difference was determined by two-way ANOVA. ns = non-significant, *p<0.05, ****p<0.0001. **H,** Western blotting of tumor lysates (3 per group) for HNF1A, BRD4, and Actin confirming knockdown of BRD4 and HNF1A downregulation/overexpression. **I,** representative immunohistochemistry staining for Ki-67 in AsPC-1 xenografts from **G**. Bar graphs represent the mean ± SEM, n=3. Statistical difference was determined by one-way ANOVA with Tukey’s multiple comparisons test; ns = non-significant, *p<0.05, **p<0.01, ***p<0.001, ****p<0.0001.

### A subset of BETi target genes is dependent on HNF1A expression

In order to gain a greater understanding of the transcriptomic contribution of HNF1A to the response of PDAC cells to BETi, we performed RNA-seq on LacZ and HNF1A overexpressing AsPC-1 cells in the presence of OTX-015 and ABBV-744, and compared the resultant gene signatures to our previously published RNA-seq of AsPC-1 cells knocked down for HNF1A^24^ (**Fig. 6A**). Using a padj.<0.05 cutoff, HNF1A knockdown resulted in 1596 downregulated and 1411 upregulated genes (**Fig. 6A, Supp. File 1**). Treatment with OTX-015 resulted in 1300 unique downregulated and 1197 unique upregulated genes, while treatment with ABBV-744 resulted in 1541 distinct downregulated and 1145 distinct upregulated genes, with 1247 downregulated (including HNF1A) and 776 upregulated genes shared by both BETi (**Fig. 6A, Supp. File 1**). Of the shared BET-regulated genes, 288 (23.1%) were also downregulated and 123 (15.9%) were upregulated by HNF1A knockdown (**Supp. File 1**). Upon re-expression of HNF1A, 1494 BETi downregulated transcripts, including 305 (20.4%) that were downregulated by HNF1A knockdown, were significantly upregulated (rescued) (**Fig. 6C**), including *HNF4A* and *EPCAM*. Collectively, we found 87/237 previously identified HNF1A-activated genes, including 23/56 associated with poor patient survival^23^, rescued by HNF1A re-expression (**Fig. 6D,E**).

**Figure 6.**
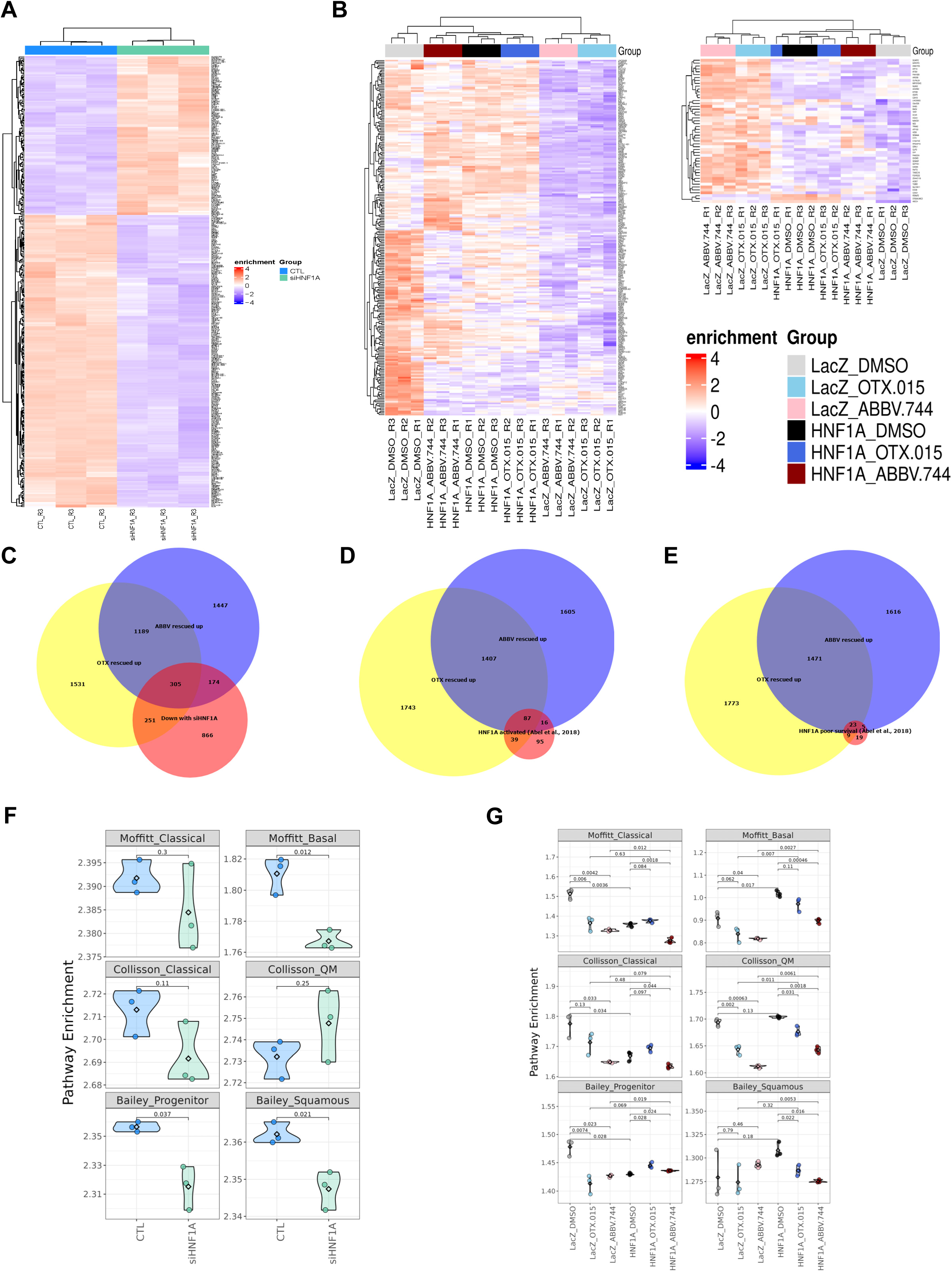
Transcriptomic analysis of HNF1A-dependent gene expression. **A,** heatmap of differentially regulated genes (DEGs) from AsPC-1 cells transfected with control or HNF1A siRNA for 72 hours; 3 biological replicates per group, (p.adj < 0.05, FC > 2). **B,** clustered heatmaps of DEGs from LacZ and HNF1A overexpressing AsPC-1 cells treated with DMSO, 0.5 µM OTX-015 or 0.1 µM ABBV-744 for 72 hours. HNF1A-rescued, BETi-downregulated transcripts are shown on the left, HNF1A-rescued, BETi-upregulated transcripts are shown on the right. 3 biological replicates per group, (p.adj < 0.05, FC > 2). **C-E,** Venn diagrams showing overlap of transcripts up-regulated by HNF1A rescue in OTX-015 and ABBV-744 treated cells with transcripts down-regulated by knockdown of HNF1A in AsPC-1 cells (**C**), previously reported HNF1A-activated genes^23^ (**D**), and HNF1A-activated genes associated with poor patient survival^23^ (**E**). **F, G,** GSVA analysis of DEGs from HNF1A knockdown (**F**) and BETi treatment/HNF1A-rescue (**G**) using previously described classical and non-classical gene sets from Collisson et al., 2011^4^, Moffitt *et al*., 2015^29^ and Bailey *et al*., 2016^28^.

Likewise,1490 BETi upregulated genes, including 260 (13.8%) that were upregulated by HNF1A knockdown were downregulated by HNF1A re-expression (Supp. Fig. 4A), including 12/45 we previously identified as HNF1A repressed genes^23^ (Supp. Fig. 4B). We had previously shown a significant overlap^24^ between HNF1A upregulated genes and genes upregulated in PDAC metastases^34^. Of these 252 overlapping transcripts, 62 (24.6%) were significantly downregulated by both BETi, with 40/62 being rescued by HNF1A re-expression, including metastasis driver *FGFR4*^24^ (**Supp. File 1**). This would indicate that BETi target metastasis associated genes and drivers in an HNF1A-dependent manner.

HNF1A is associated with the classical subtype of PDAC^28, 35^ and has been proposed to serve as a barrier to transdifferentiation into non-classical subtypes^8^. To determine whether manipulation of HNF1A and/or BETi drive subtype switching in PDAC, we performed gene set variation analysis (GSVA) using well-established PDAC subtype gene sets^4, 28, 29^. While the classical signatures defined by Collisson *et. al.*, and Moffitt *et. al.*, were not significantly altered by HNF1A knockdown, we did observe a downregulation of the analogous Bailey *et. al.*, progenitor subtype signature. Interestingly, the knockdown of HNF1A also downregulated the Moffitt *et. al.*, basal subtype and Bailey *et. al.*, squamous subtype signatures without significantly altering the Collisson quasimesenchymal (QM) subtype signature (**Fig. 6F**), indicating that HNF1A positively regulates genes associated with multiple subtypes, similar to our previous work^23^. Likewise, we did not observe an induction of the non-classical signatures with BETi treatment, but rather a suppression of these signature genes in most cases and in a similar manner to classical/progenitor genes (**Fig. 6G**). Overexpression of HNF1A significantly downregulated the classical/progenitor signatures upregulated the Collison *et. al.*, QM signature (**Fig. 6G**), indicating that HNF1A’s role in subtype determination is likely more complicated than previously suggested^8^.

### Receptor tyrosine kinase (RTK) signaling is coordinated by HNF1A downstream of BETi

In order to determine the mechanism by which HNF1A re-expression rescues the antiproliferative effects of BETi, we performed gene set enrichment analysis (GSEA) using our BETi/HNF1A rescue RNA-seq results. As expected, numerous pathways were altered by both BETi; however, only a small subset of pathways showed rescue by HNF1A re-expression (49 pathways rescued up, 13 pathways rescued down) (Supp. Fig 4C). Among the 49 pathways that were downregulated by BETi and rescued by HNF1A re-expression, six involved components of EGFR/ERBB-signaling (PID_ERBB_NETWORK_PATHWAY, GOBP_REGULATION_OF_EPIDERMAL_GROWTH_FACTOR_ACTIVATED_RECEPTOR_ACT IVITY, GOBP_EPIDERMAL_CELL_DIFFERENTIATION, GOBP_KERATINOCYTE_PROLIFERATION, GOBP_POSITIVE_REGULATION_OF_ERBB_SIGNALING_PATHWAY, and GOBP_REGULATION_OF_KERATINOCYTE_PROLIFERATION) (Supp. Fig 4C). In examining our HNF1A knockdown and BETi rescue RNA-seq data, we observed multiple components of EGFR/ERBB-signaling significantly altered, including mRNA levels of receptors (*EGFR* and *ERBB3*), ligands (*AREG*, *EREG*, *TGFA*, and *HBEGF*), and regulator proteins (*TRIB3* and *ERRFI1*) showing downregulation by HNF1A knockdown and BETi and rescued expression by HNF1A restoration (**Fig. 7A, B**). Consistent with our mRNA data, we observed decreased levels of total and/or phosphorylated EGFR and ERBB3 across multiple PDAC cell lines in response to BETi treatment, which was in turn rescued by re-expression of HNF1A (**Fig. 7C**, Supp. Fig. 5A). Similar results were observed with direct knockdown of BRD4 and HNF1A re-expression (Supp. Fig. 5B), indicating that EGFR/ERBB3-signaling are both responsive to BETi/BRD4-ablation and are HNF1A-dependent. To determine whether restoration of EGFR/ERBB3-signaling was responsible for the rescue phenotype imparted by HNF1A restoration, we targeted EGFR activity with the EGFR-specific inhibitor, erlotinib, and pan-ERBB inhibitor, afatinib. Both inhibitors reduced the phosphorylation of EGFR and ERBB3 (**Fig. 7D**, Supp. Fig. 5C) and abolished the protective effects of HNF1A re-expression in the presence of BETi (**Fig. 7E**, Supp. Fig. 5D). While single-agent activity of both inhibitors was observed, no baseline differences in sensitivity due to HNF1A re-expression was seen, supporting that EGFR/ERBB3-signaling functions downstream of HNF1A. To determine whether HNF1A was more broadly associated with EGFR/ERBB3-signaling components in PDAC, we mined the Moffitt *et. al.*, and Bailey *et. al.*, datasets for correlations between *HNF1A* mRNA and expression levels of *EGFR*, *ERBB3*, *AREG*, *EREG*, *TGFA*, *HBEGF*, *TRIB3* and *ERRFI1*. Of these genes, *HNF1A* mRNA was correlated with *ERBB3* mRNA across both datasets (**Fig. 7F, G**), while *TRIB3* and *ERRFI1* mRNA only in the Moffitt *et. al.*, dataset (Supp. Fig. 6A, B). Examining our own HNF1A ChIP-seq datasets^23, 24^, we identified a strong enrichment peak for HNF1A approximately −77kbp to the transcriptional start of ERBB3 in both UM5 and UM15, indicating possible distal regulation of ERBB3 by HNF1A in PDAC (Supp. Fig. 6C).

**Figure 7.**
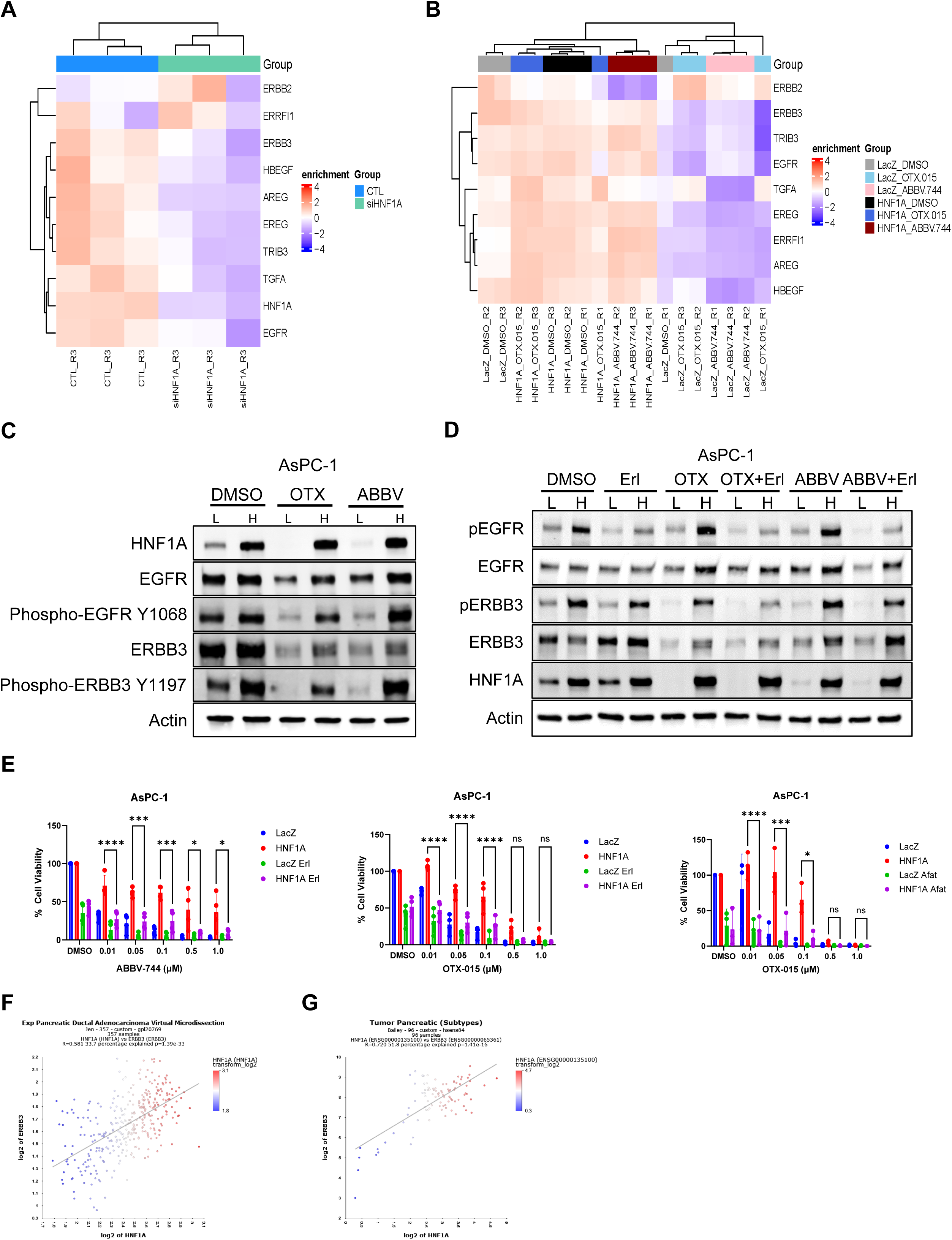
HNF1A re-expression drives EGFR/ERBB3-signaling. **A, B,** heatmap of differentially expressed genes in the EGFR/ERBB3-signaling pathway from RNA-seq of AsPC-1 cells transfected with HNF1A siRNA (**A**) or overexpressing LacZ and HNF1A and treated with the indicated BETi for 72 hours (**B**). **C,** Western blotting for phospho-EGFR Y1068, EGFR, phospho-ERBB3 Y1197, ERBB3, HNF1A, and Actin from LacZ (L) and HNF1A (H) overexpressing AsPC-1 cells treated with DMSO, 0.5 µM OTX-015, or 0.1 µM ABBV-744 for 72 hours. **D,** Western blotting for phospho-EGFR Y1068, EGFR, phospho-ERBB3 Y1197, ERBB3, HNF1A, and Actin from LacZ (L) and HNF1A (H) overexpressing AsPC-1 cells treated with DMSO, 1 µM erlotinib, 0.5 µM OTX-015, 0.1 µM ABBV-744, or combinations of erlotinib and either OTX-015 or ABBV-744 for 72 hours. **E,** dose curves colony formation experiments for LacZ, and HNF1A overexpressing AsPC-1 cells treated with DMSO, 1 µM erlotinib, 0.5 µM OTX-015, 0.1 µM ABBV-744, or combinations of erlotinib and either OTX-015 or ABBV-744 (left panels) or BETi in combination with 1 µM afatinib (right panel) for two weeks. Bar graphs represent the mean ± SEM, n=3. Statistical difference was determined by one-way ANOVA with Tukey’s multiple comparisons test; ns = non-significant, *p<0.05, ***p<0.001, ****p<0.0001. **F, G,** correlation of HNF1A and ERBB3 mRNA expressions from Moffitt *et al*., 2015^29^ (**F**) and Bailey *et al*., 2016^28^ (**G**).

### HNF1A, BRD4, and RTK expression are correlated in PDAC patients

In the PDAC cell models examined in this study we observed an epistatic hierarchy wherein BETi target BRD4, which directly regulates HNF1A, which either directly or indirectly drives EGFR/ERBB3-signaling to promote PDAC proliferation and stemness. To examine whether this hierarchy exists more broadly in PDAC patients, we performed multispectral immunofluorescence on a tissue microarray (TMA) consisting of 220 PDAC patient cores^26^.

Using pan-cytokeratin (AE1/AE3) to label PDAC cells, we co-stained cores for HNF1A, BRD4, ERBB3, phospho-ERBB3, and established HNF1A target and RTK FGFR4^24^ (**Fig. 8A, B**). Using both percent positivity and staining intensity, we found that HNF1A and BRD4 were the most strongly correlated of the markers tested (**Fig. 8C-E**), with these markers showing significant correlations when measuring percent positivity (**Fig. 8D**). Interestingly, when examining the correlation coefficients (Pearson R) for staining intensities for each core, we found the R values for FGFR4 and ERBB3 to be the strongest of all pairwise comparisons (**Fig. 8E**). As both of these RTKs are part of a metastatic signature in PDAC (**Supp. File 1**)^24^, these findings lend credence to the possibility that HNF1A orchestrates PDAC malignancy, at least in part, through upregulation of RTK-signaling, and that this axis is vulnerable to BETi. Finally, we examined whether the staining intensity of our markers was correlated with patient outcome in the 115 patient samples with available survival data. Of these markers, only ERBB3 was significantly correlated with survival, with higher ERBB3 staining intensity associated with better survival (Supp. Fig. 6D). Overall, these findings suggest that the BRD4-HNF1A-RTK regulatory axis may have both therapeutic and prognostic value in PDAC.

**Figure 8.**
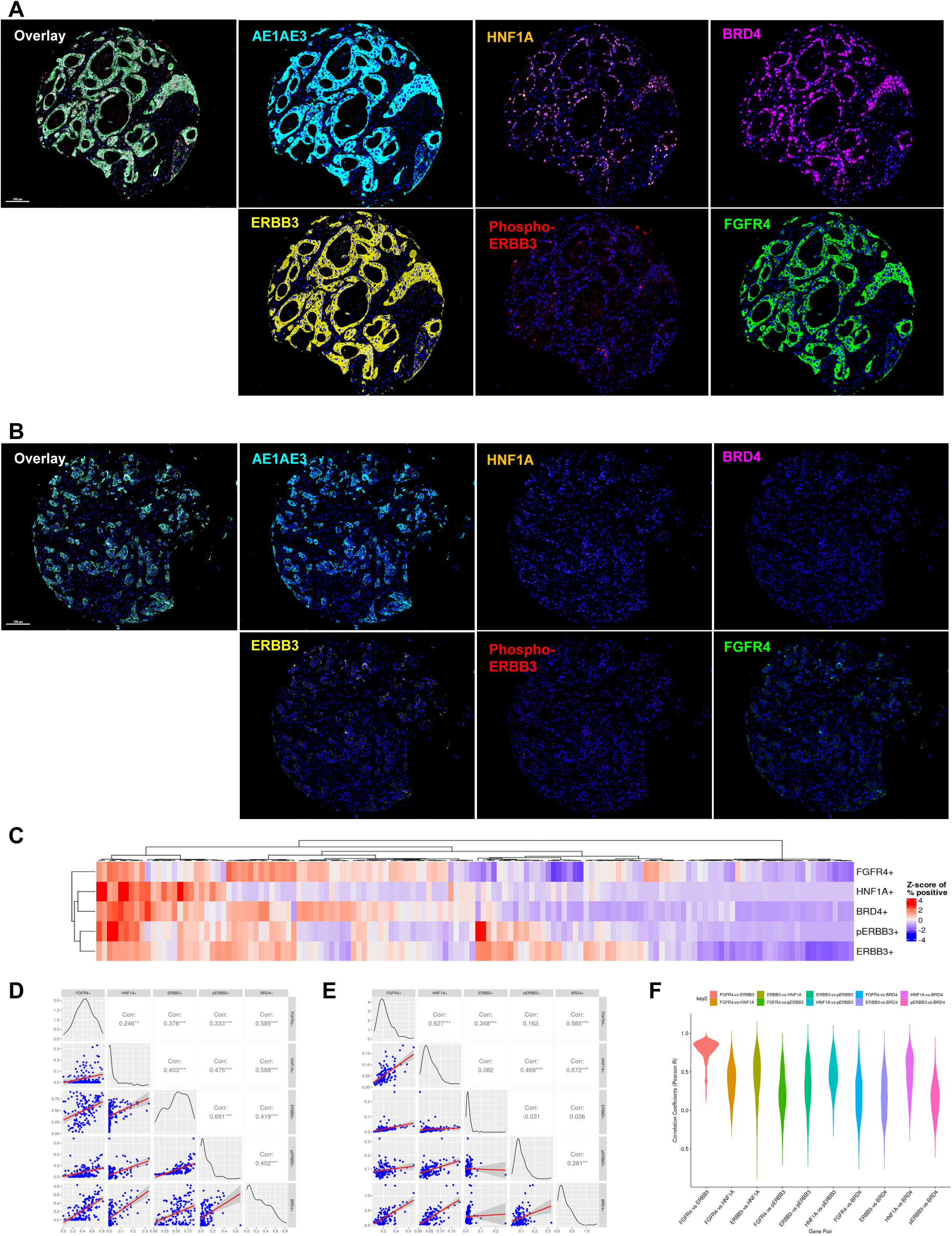
Co-expression of HNF1A, BRD4, and RTKs in PDAC patient samples. **A, B,** multispectral immunofluorescence of PDAC patient cores staining for pan-cytokeratin (AE1AE3), HNF1A, BRD4, ERBB3, phospho-ERBB3 Y1289, and FGFR4. A core demonstrating strong co-staining for all markers is represented in (**A**), while a core demonstrating only strong staining for pan-cytokeratin is represented in (**B**). **C,** ranked heatmap showing correlation of HNF1A, BRD4, ERBB3, phospho-ERBB3, and FGFR4 positivity across 220 patient cores. **D, E,** correlation plots for markers based on percent positivity (**D**) and average staining intensity (**E**). **F,** violin plot of the R values for each core comparing marker staining intensity. Pairwise comparisons are indicated above.

## DISCUSSION

In this study, we demonstrated that the oncogenic transcription factor HNF1A is directly regulated by BRD4, and therefore sensitive to BETi, and that this suppression of HNF1A expression is a major contributor to the antineoplastic activity of BETi in HNF1A-expressing PDAC cells. These findings uncover a novel mechanism of BETi activity in PDAC and highlight HNF1A as a potential therapeutic target.

Pharmacological inhibition of epigenetic writers (e.g., HATs), erasers (e.g., HDACs), and readers (e.g., BET proteins) have been widely pursued in the treatment of many cancers and have been explored in clinical trials for PDAC^36^. However, in the case of BETi, enrollment of PDAC patients in clinical trials has been limited, making the efficacy of BETi challenging to assess. Nonetheless, larger PDAC-focused clinical trials examining the efficacy of BETi would likely be met with a number of challenges. One foreseeable limitation would be the evaluation of effective BETi delivery and on-target activity, particularly as the tumor microenvironment may present a barrier to drug delivery^37^. While MYC has long been regarded as the most notable target of BETi and may serve as a biomarker for BETi efficacy, the responsiveness of MYC expression to BETi in PDAC and other cancers is variable^15, 38^, and further supported by this study. Here, we found HNF1A expression exquisitely sensitive to BETi, even at doses and in cell lines in which MYC was not responsive. It is therefore feasible that the expression of HNF1A in patient tumors may serve as a biomarker for BETi activity as well as a critical downstream target of BRD4. Another limitation of BETi in cancer therapeutics is on-target toxicities caused by the disruption of BET-dependent gene transcription, and as such, bromodomain-selective inhibitors with more restricted effects have been proposed as safer alternatives to pan-BETi^39, 40^. ABBV-744 was initially demonstrated to have potent antiproliferative activity against prostate cancer while exhibiting a more modest effect on gene expression than pan-BETi^39^. In our hands, ABBV-744 and OTX-015 exhibited very distinct effects on gene expression, supporting that the activity of BD2-selective BETi and pan-BETi are indeed distinct in PDAC cells; however, we were able to demonstrate that both classes of BETi disrupt HNF1A-dependent gene expression and inhibit PDAC cell growth and stem-like properties in an HNF1A-dependent manner. This supports the idea that multiple classes of BETi, including future generations of improved BETi or BRD4-targeting modalities may be utilized to target HNF1A in PDAC or other HNF1A-driven cancer types. Alternatively, direct inhibition of HNF1A activity/expression (e.g., PROTAC or RNAi) may serve as a strategy to cripple its transcriptional network in PDAC cells without widespread transcriptional disruption seen with inhibition of common epigenetic/transcriptional regulators.

The role(s) of HNF1A in cancer, particularly PDAC, remain controversial. HNF1A expression has been shown to mark and possibly contribute to early onset resistance to androgen-ablation in prostate cancer^22^, as well as drive super-enhancer remodeling and liver metastasis in colorectal cancer^20^. Early genome-wide association (GWA) studies in PDAC identified multiple risk variant single nucleotide polymorphisms (SNPs) of the HNF1A locus^41–43^, although no studies to date have ascribed a function to any of these risk variants that would support gain or loss of HNF1A function or expression. Supporting an oncogenic role for HNF1A in PDAC, our group and others have demonstrated that HNF1A promotes metastasis^44^, maintains cancer stem cell function^23^, and is necessary for tumor growth in HNF1A-expressing PDAC cells^23, 45^. By contrast, Hoskins *et al*., was the first to suggest HNF1A as a putative tumor suppressor in PDAC, observing that the HNF1A transcriptional network was dysregulated in PDAC^46^, a phenomenon that can now be attributed to the different molecular subtypes of PDAC and their unique transcriptional regulators^4, 28, 29, 35, 47^. Additionally, overexpression of HNF1A in non-classical PDAC cells (Panc-1 and MiaPaCa-2) resulted in growth arrest and apoptosis, possibly due to super-physiological levels of HNF1A expression or incompatibility of HNF1A with endogenous transcriptional programs. Additionally, the requirement for HNF1A in HNF1A-expressing cells was not examined. Of note, we did not observe anti-proliferative effects from overexpression of HNF1A from the weaker UbC promoter in either Panc-1 or MiaPaCa-2 (data not shown), suggesting that the dose of overexpression may affect HNF1A’s proliferative/anti-proliferative activities. In addition to being proposed as a tumor suppressor, it has also been suggested that HNF1A, in concert with GATA6 and HNF4A^8^ or KDM6A^48^, might serve as transcriptional roadblock to the non-classical subtype(s) of PDAC, potentially complicating the implementation of HNF1A-targeting therapies. While we found that BETi and short-term knockdown of HNF1A diminished the classical gene signature in AsPC-1 cells, we did not observe a significant increase in non-classical gene expression. We also observed positive regulation of some non-classical signature genes, such as *AREG*, by HNF1A, indicating that HNF1A does not solely regulate classical signature genes. These observations are consistent with our previous long-term knockdown studies in PDX lines^23^, indicating that loss of HNF1A alone is insufficient to transdifferentiate classical PDAC to non-classical PDAC. Nonetheless, non-classical PDAC cells have been shown to be sensitive to BETi^16^, presumably through disruption of their unique master transcription factors like MYC, RUNX3, and TP63^9, 16^, and taken together with this study, support the use of BETi to overcome critical transcriptional networks.

Key among the findings of this study are the connections between HNF1A, BETi, and EGFR/ERBB3-signaling. While resistance to BETi mediated through EGFR-signaling has been reported in other cancers^49^, the role of this pathway in response to BETi in PDAC has not been explored, nor has its regulation by HNF1A. Recently, Rao *et al*. reported that high expression of HNF1A could be used to stratify patient responses to a combination of gemcitabine and erlotinib, with HNF1A positivity correlating with better survival. While this study did not explore whether HNF1A status correlated with expression or phosphorylation of EGFR or ERBB3, it is in line with our observation that ERBB3-high patients within our cohort had better outcomes. This finding, however, disagrees with early observations that high ERBB3 predicted poorer survival in patients with resected PDAC^50^, though this may reflect differences in how ERBB3, and perhaps HNF1A, impact treatment responses. It is worth noting that ERBB3 was among the classical PDAC signature genes initially described by Collisson *et al*., who also demonstrated that classical PDAC cells were more sensitive to erlotinib than quasi-mesenchymal PDAC cells^4^.

While it is unclear whether HNF1A directly regulates ERBB3, we previously identified it as an HNF1A-regulated gene and identified an HNF1A binding peak ∼77 kbp upstream of the ERBB3 transcriptional start site^23^. Overall, these findings would suggest that HNF1A, rather than functioning as a *bona fide* tumor suppressor, transcriptionally controls signaling networks (such as RTKs) that can be exploited for therapeutic gain.

A perplexing finding of this study is the interplay between BETi, HNF1A, and MYC. Many early studies on BETi in leukemias suggest that their antineoplastic activity stem from the transcriptional inhibition of MYC^12, 17–19^, although their activity in PDAC PDXs has been demonstrated to not strictly correspond to inhibition of MYC^15^. Consistent with the latter, we found that BETi variably affected MYC expression in PDAC cell and PDX lines, and generally less effectively than their inhibition of HNF1A expression. Moreover, ectopic expression of MYC, in contrast to ectopic HNF1A, did not rescue the antiproliferative effects of BETi, indicating that inhibition of MYC alone is not the determinant of BETi activity in HNF1A-expressing PDAC cells. However, HNF1A re-expression also rescued MYC protein (Fig. 2A) and mRNA levels (Fig. 5B, Supplemental File 2), suggesting that HNF1A may control MYC expression independently of BRD4, possibly bolstering HNF1A oncogenic activity in conjunction with RTK upregulation.

Future studies may be warranted to better understand the potential crosstalk between MYC and HNF1A in PDAC and other HNF1A-expressing cancers.

In summary, this study highlights a novel role for HNF1A in determining the response of HNF1A-expressing PDAC to BETi and may in part explain the association between HNF1A-positive/classical subtype PDAC and responsiveness to EGFR-inhibitors. While these findings have great implications for stratifying the use of targeted therapies like BETi and EGFR-inhibitors in classical PDAC, it remains unclear whether this paradigm would apply to non-classical PDAC, or what driver transcription factors might portend response in this subtype. A better understanding of the critical targets of BETi in HNF1A-negative/nonclassical is needed not only as potential biomarkers for response to BETi, but also to identify other potential vulnerabilities, like the EGFR pathway in HNF1A-positive PDAC. Future studies should also explore whether HNF1A expression, as well as its downregulation by BETi, can be used to stratify patients and predict response to BETi in the clinic.

## AUTHORS’ CONTRIBUTIONS

**K.S. Humphrey:** Conceptualization, data curation, formal analysis, validation, investigation, methodology, and writing–original draft. **K.J. Crawford:** Data curation, formal analysis, validation, investigation, writing–original draft, and editing. **B. Muppavarapu:** Data curation, formal analysis, validation, investigation, funding acquisition and methodology. **M.M. Mayberry:** Data curation, formal analysis, validation, investigation, and methodology. **W. Morris:** Data curation, formal analysis, validation, investigation, and methodology. **E. Torres:** Data curation, formal analysis, validation, investigation, and methodology. **M.D. Long:** Data curation, software, and formal analysis. **J. Wang:** Data curation, software, methodology, and formal analysis. **E.S. Knudsen:** Conceptualization, funding acquisition, resources, and editing. **A.K. Witkiewicz:** Conceptualization, funding acquisition, resources, and editing. **E.V. Abel:** Conceptualization, data curation, formal analysis, validation, investigation, supervision, funding acquisition, methodology, project administration, and writing–original draft.

## ACKNOWLEDGMENTS

We thank all the members of the laboratory group and colleagues in the discussion and preparation of this manuscript, especially the labs of Drs. Howard Crawford (Henry Ford Health), Michael Feigin (RPCCC) and Anna Bianchi-Smiraglia (RPCCC). We would like to thank Sidney Mahan (RPCCC) and Hanna Rosenheck (RPCCC) for helping to optimize and perform multispectral immunofluorescence, and Robert Kyne (RPCCC) for assisting with animal studies. This work was supported by Roswell Park Comprehensive Cancer Center and National Cancer Institute (NCI) grant, P30CA016056; NCI grant R37CA275961, Pancreatic Cancer Action Network-AACR Pathway to Leadership Grant (16-70-25-ABEL), the SAS Foundation for Cancer Research Grant, the Roswell Park Alliance Foundation, and the Hirshberg Foundation for Pancreatic Cancer Research Seed Grant (to EVA); the Mark Diamond Research Fund grant (to KSH and BM), and NCI grants R01CA267467 and R01CA211878 (to EK and AW). This work was supported by Roswell Park Comprehensive Cancer Center and National Cancer Institute (NCI) grant, P30CA016056. The content is solely the responsibility of the authors and does not necessarily represent the official view of the National Institutes of Health.

---

**Supplemental Figure 1.**
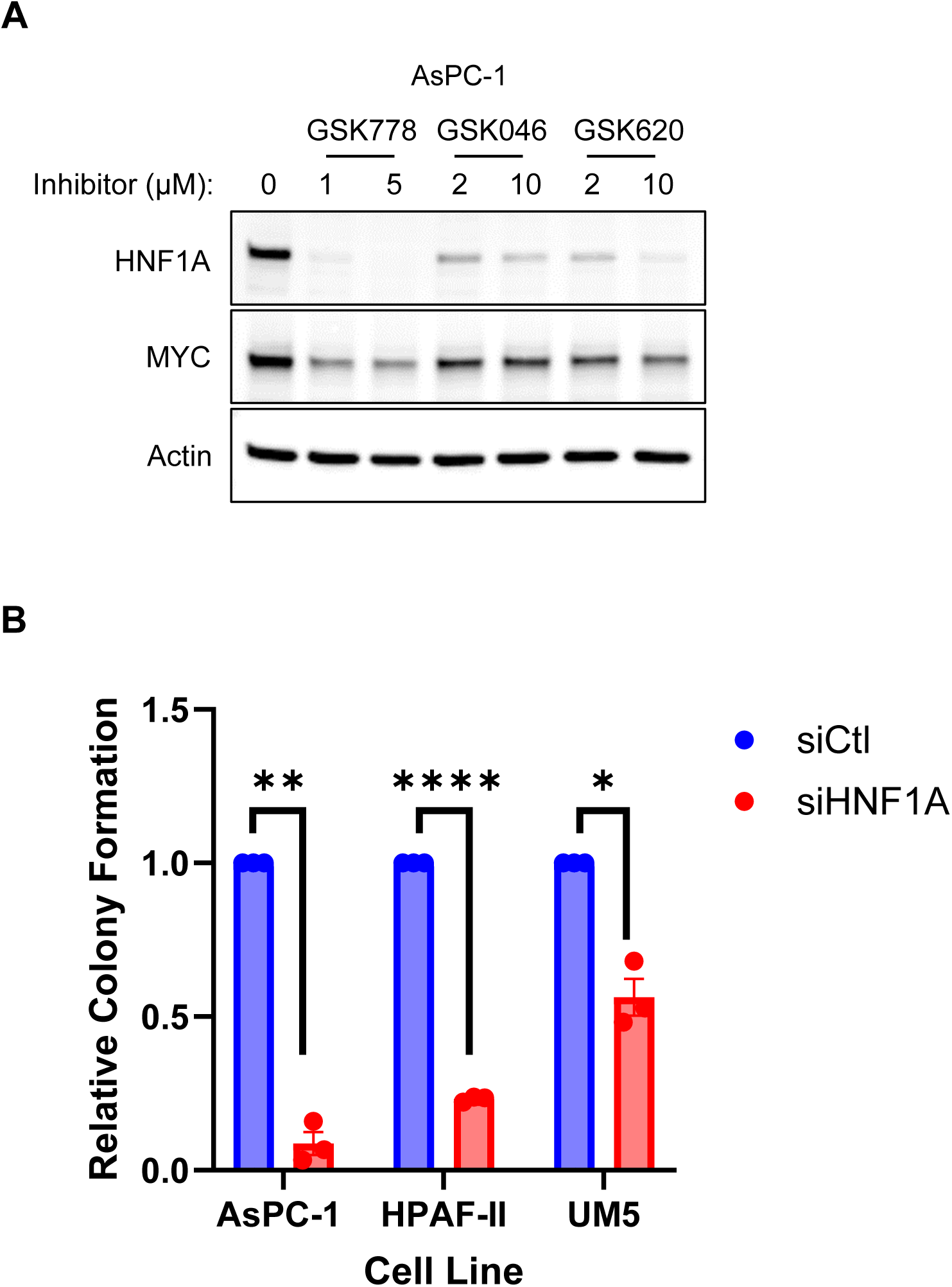
Inhibition of HNF1A expression by BD-selective BET-inhibitors and effects of HNF1A loss on colony formation. **A,** Western blotting of HNF1A, MYC, and Actin for AsPC-1 cells were treated with the indicated concentrations of GSK778 (BD1-selective), GSK046 (BD2-selective), or GSK620 (BD2-selective) for 72 hours. **B,** Colony formation of AsPC-1, HPAF-II, and UM5 following knockdown of HNF1A. Bar graphs represent the mean ± SEM, n=3. Statistical difference was determined by two-sided Student t-test with Welch’s correction; *p<0.05, **p<0.01, ****p<0.0001.

**Supplemental Figure 2.**
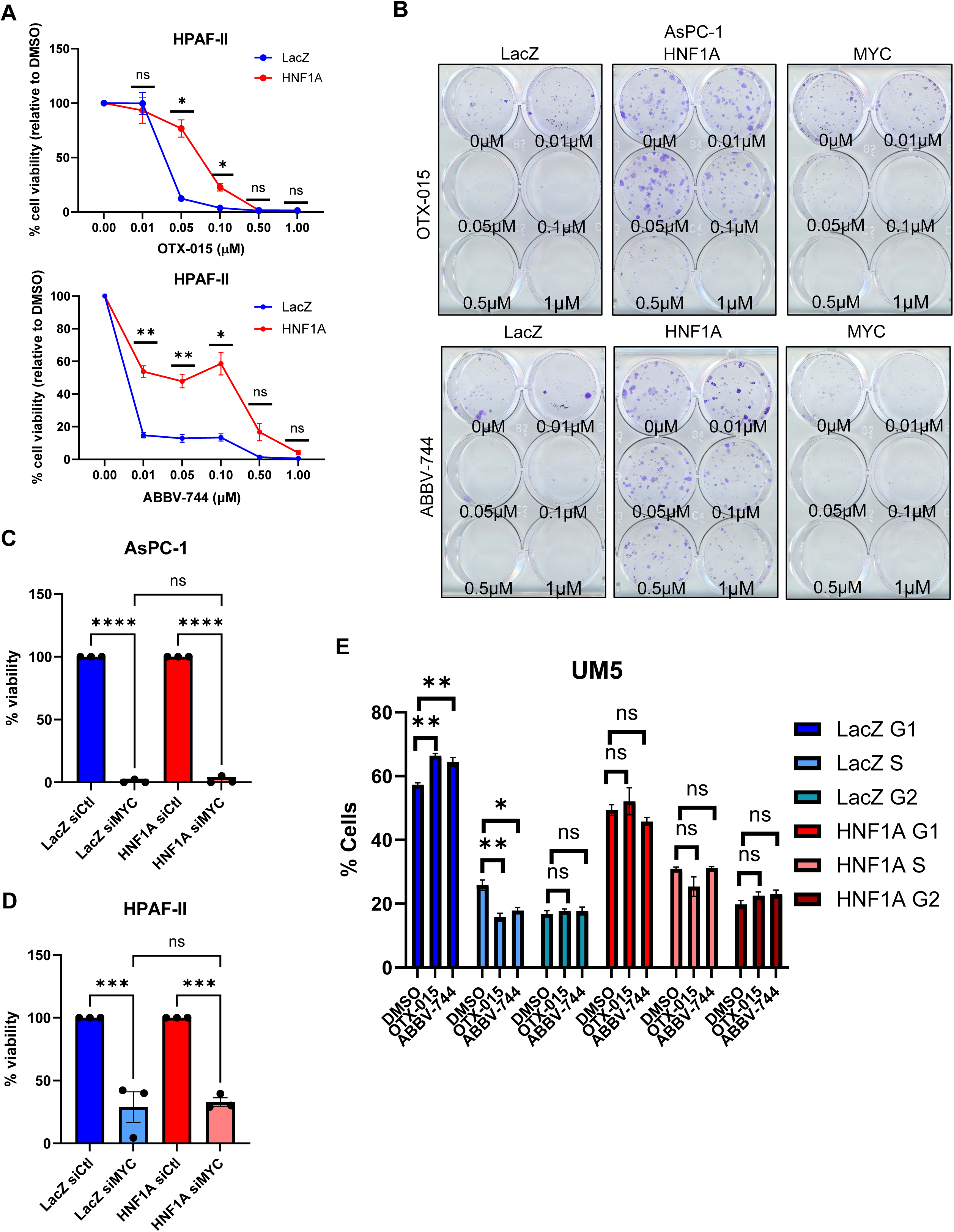
HNF1A overexpression protects against BETi. **A,** dose curves colony formation experiments for LacZ and HNF1A overexpressing HPAF-II cells. Cells were plated at 200 cells per well and treated with the indicated concentrations of OTX-015 or ABBV-744 for 2 weeks. Resultant colonies were quantitated by AlamarBlue viability assay. Data points represent the mean ± SEM, n=3. Statistical difference was determined by one-way ANOVA with Tukey’s multiple comparisons test; ns = non-significant, *p<0.05, **p<0.01. **B,** representative LacZ, HNF1A, and MYC overexpressing AsPC-1 colonies after 2 weeks of BETi treatment. **C, D,** colony viability of AsPC-1 (**C**) and HPAF-II (**D**) LacZ and HNF1A overexpressing cells transfected with MYC siRNA for 2 weeks. **E,** quantitation of propidium iodide staining of LacZ and HNF1A overexpressing UM5 cells treated with 0.5 µM OTX-015 or 0.1 µM ABBV-744 for 72 hours. Relative percentages of G0/G1, S, and G2/M phase cells are shown. Statistical difference was determined by one-way ANOVA with Tukey’s multiple comparisons test; ns = non-significant, *p<0.05, **p<0.01, ***p<0.001, ****p<0.0001.

**Supplemental Figure 3.**
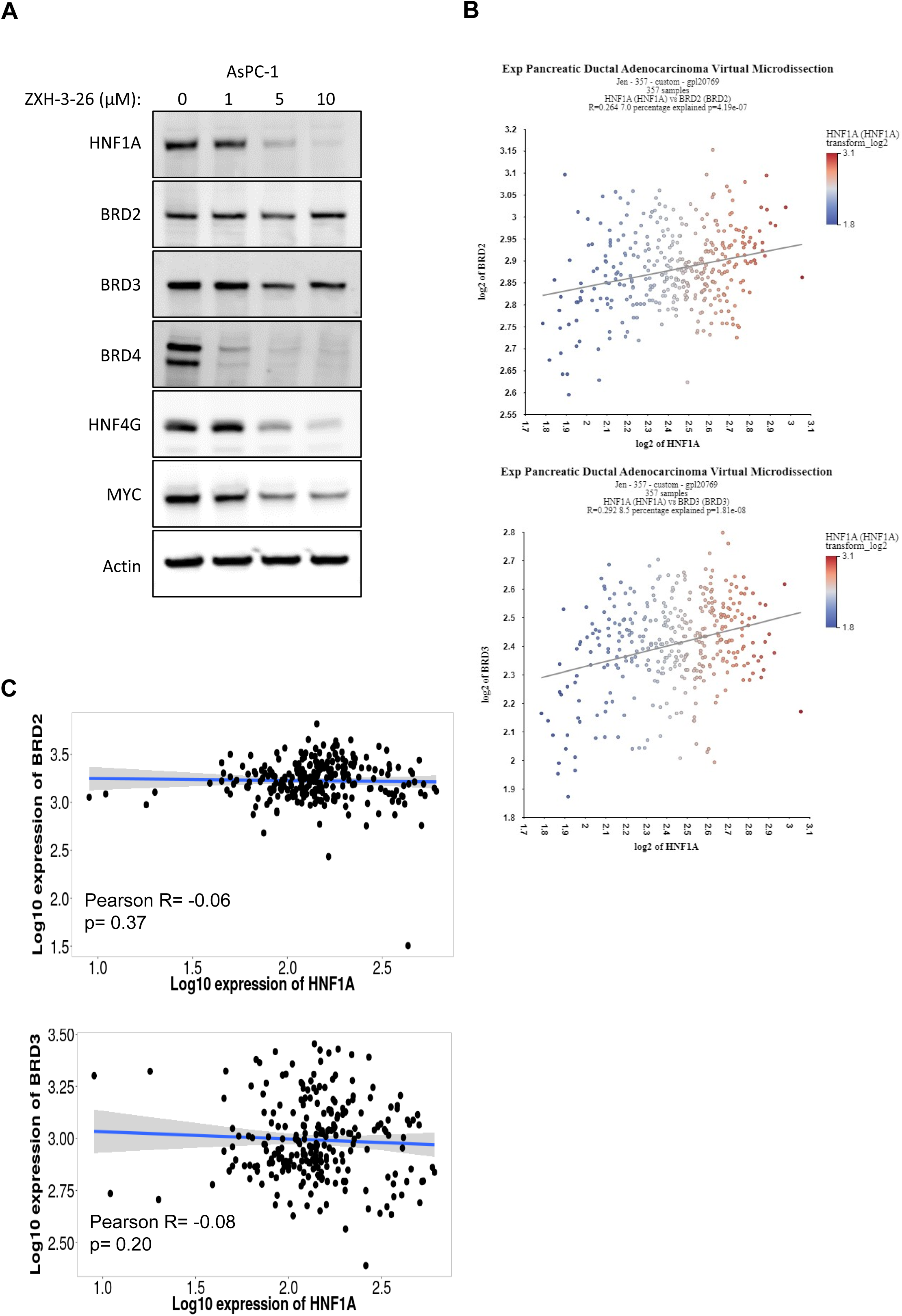
Association of HNF1A with BET-family proteins in PDAC. **A,** Western blotting of HNF1A, BRD2, BRD3, BRD4, HNF4G, MYC, and Actin in AsPC-1 cells treated with the indicated concentrations of BRD4-selective degrader ZXH-3-26 for 72 hours. **B,** correlations of HNF1A mRNA expression and BRD2 and BRD3 mRNA expressions from Moffitt *et al*., 2015^29^. **C,** correlations of HNF1A mRNA expression and BRD2 and BRD3 mRNA expressions from PDAC tumors from TNMplot.com^33^.

**Supplemental Figure 4.**
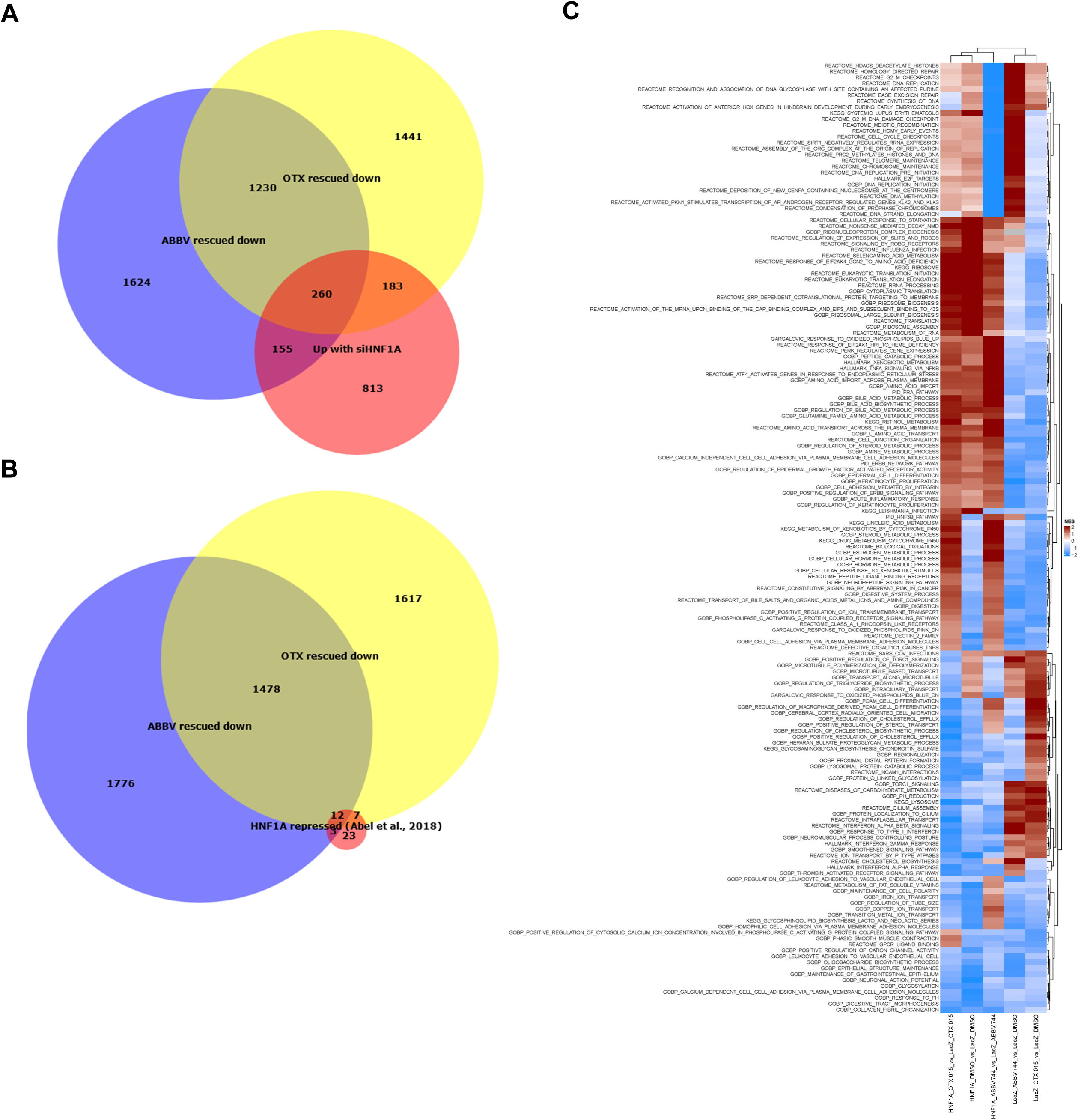
Transcriptomic and pathway analysis of BETi-responsive/HNF1A-dependent genes. **A, B,** Venn diagrams showing overlap of transcripts downregulated by HNF1A rescue in OTX-015 and ABBV-744 treated cells with transcripts upregulated by knockdown of HNF1A in AsPC-1 cells (**A**) and previously reported HNF1A-repressed genes^23^ (**B**). **C,** clustered GSEA heatmap for pathways differentially regulated by OTX-015 or ABBV-744 treatment and HNF1A rescue in AsPC-1 cells.

**Supplemental Figure 5.**
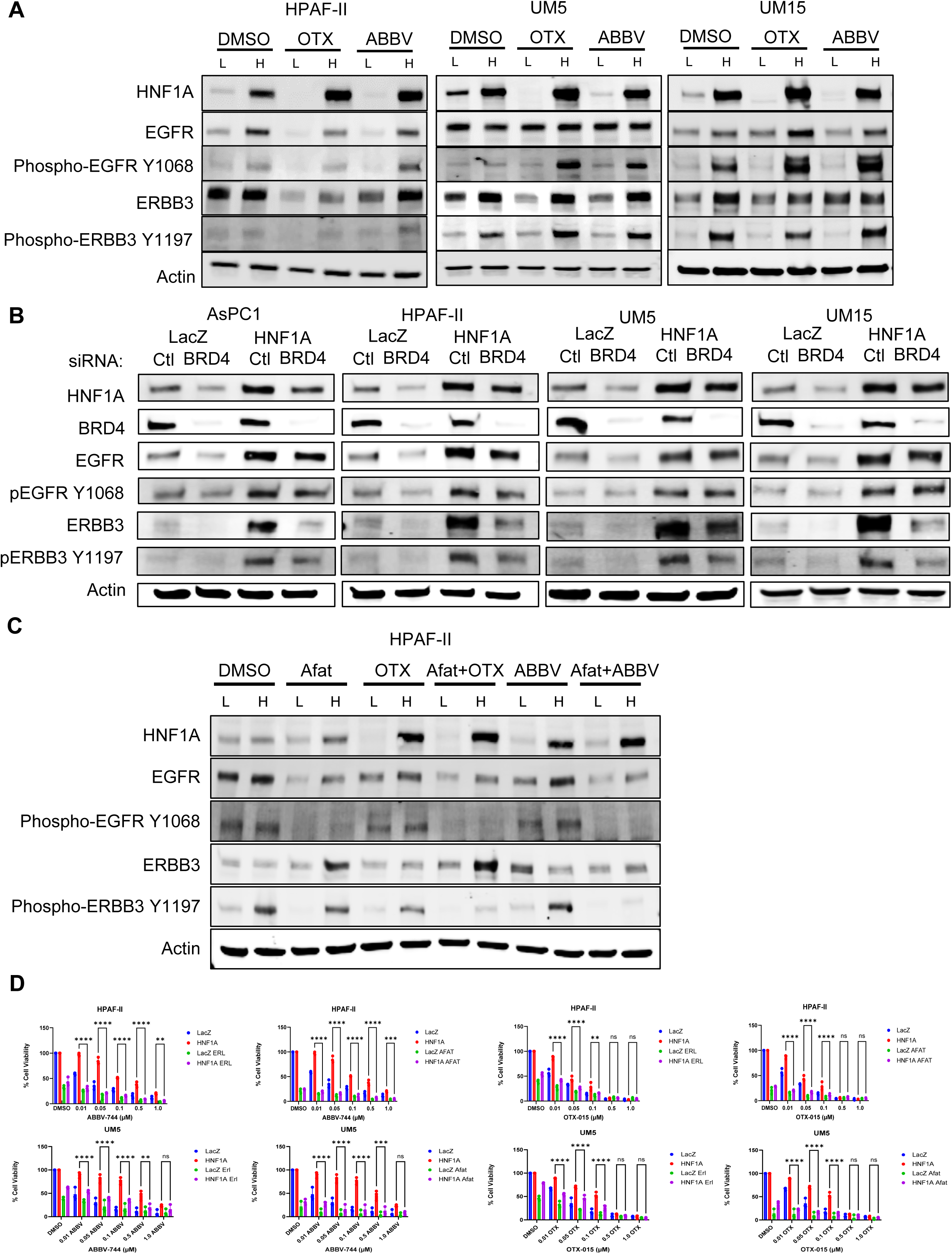
Regulation of EGFR/ERBB3-signaling by HNF1A, BETi, and BRD4. **A,** Western blotting for phospho-EGFR Y1068, EGFR, phospho-ERBB3 Y1197, ERBB3, HNF1A, and Actin from LacZ (L) and HNF1A (H) overexpressing HPAF-II, UM5, and UM15 cells treated with DMSO, 0.5 µM OTX-015, or 0.1 µM ABBV-744 for 72 hours. B, Western blotting for phospho-EGFR Y1068, EGFR, phospho-ERBB3 Y1197, ERBB3, HNF1A, and Actin from LacZ and HNF1A overexpressing AsPC-1, HPAF-II, UM5, and UM15 cells knocked down for BRD4. **C,** Western blotting for phospho-EGFR Y1068, EGFR, phospho-ERBB3 Y1197, ERBB3, HNF1A, and Actin from LacZ (L) and HNF1A (H) overexpressing HPAF-II cells treated with DMSO, 1 µM afatinib, 0.5 µM OTX-015, 0.1 µM ABBV-744, or combinations of afatinib and either OTX-015 or ABBV-744 for 72 hours. **D,** dose curves colony formation experiments for LacZ, and HNF1A overexpressing HPAF-II and UM5 cells treated with DMSO, 1 µM erlotinib, 0.5 µM OTX-015, 0.1 µM ABBV-744, or combinations of erlotinib and either OTX-015 or ABBV-744 (left panels) or BETi in combination with 1 µM afatinib (right panel) for two weeks. Bar graphs represent the mean ± SEM, n=3. Statistical difference was determined by one-way ANOVA with Tukey’s multiple comparisons test; ns = non-significant, *p<0.05, ***p<0.001, ****p<0.0001.

**Supplemental Figure 6.**
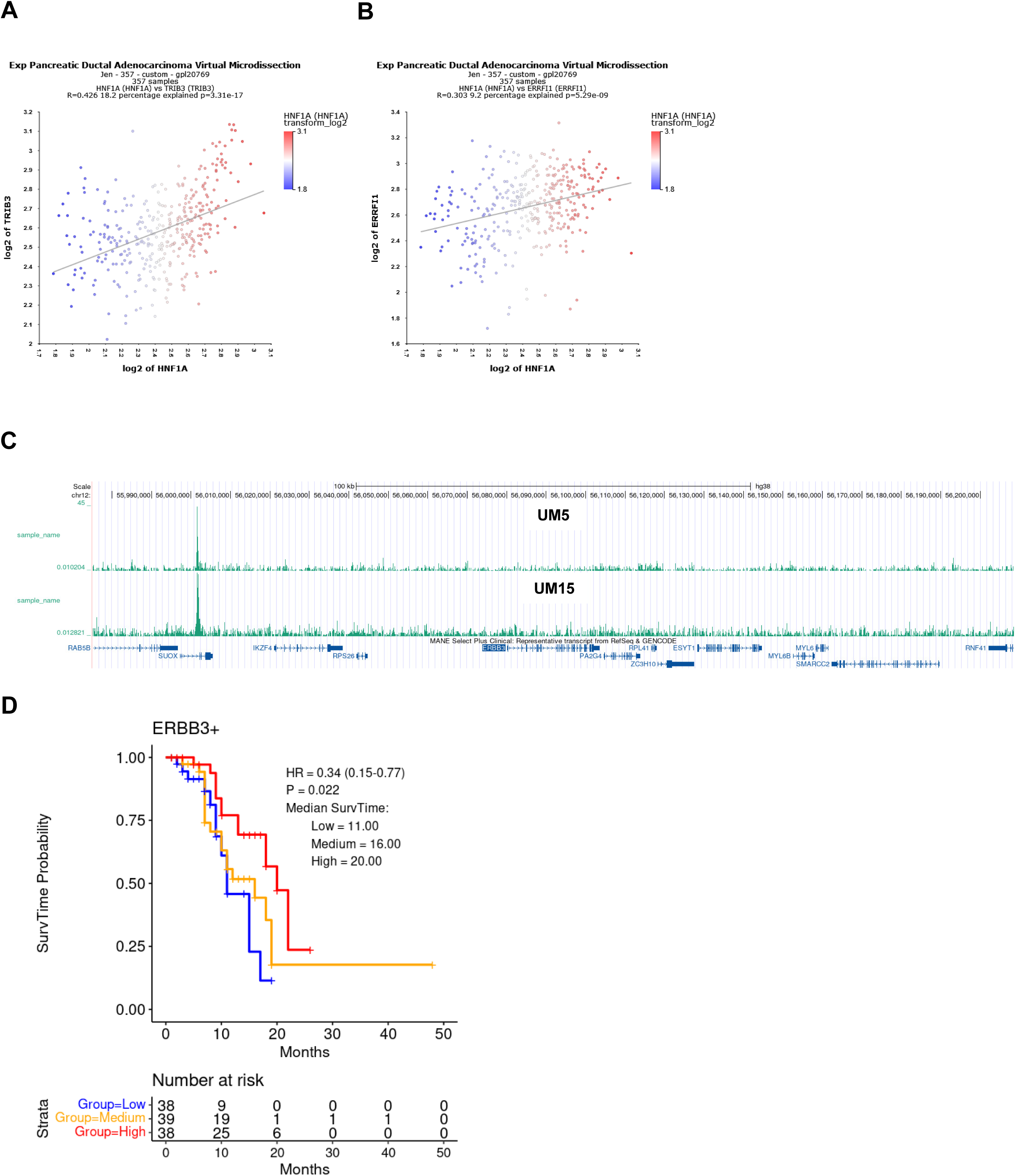
Association of HNF1A with EGFR/ERBB3-signaling components. **A, B,** Correlation of *TRIB3* (**A**) and *ERRFI1* (**B**) mRNA expressions with *HNF1A* mRNA expression from Moffitt *et al*., 2015^29^. **C,** UCSC Genome Browser HNF1A ChIP-seq tracks from UM5 and UM15 cells showing the ERBB3 locus and adjacent gene bodies. **D,** Kaplan-Meier plot for PDAC patient survival correlated with ERBB3 staining intensity. Patients were stratified by low, medium, and high expression of ERBB3. N=115 patients.

